# Changes in enclosure use and basking behaviour associated with pair housing in Tokay geckos (*Gekko gecko*)

**DOI:** 10.1101/2024.01.08.574605

**Authors:** Birgit Szabo

## Abstract

Due to often insufficient information reptiles suffer welfare issues and increased mortality in captivity. In particular, the impact of the social environment remains poorly understood, despite evidence suggesting its’ importance for welfare in a wide range social animals. The current study investigated how pair housing changes enclosure use, basking and hiding behaviour in tokay geckos (*Gekko gecko*). While the captive conditions and husbandry procedures employed in this study align with existing literature recommendations, they have not been previously evaluated for their suitability for this particular species. The results show that, when socially deprived, lizards were more likely to move and hide before feeding. Furthermore, males were more likely to be found at the front than females during pair housing but not during single housing. Finally, contrary to single housing, enclosure temperature had no effect on the probability to move and hide behind a shelter during pair housing. Consistently, however, lizards were more likely to bask after feeding across housing conditions and females were more likely to bask before their first clutch. Together, pair housing decreases movement and hiding in relation to human presence (feeding) which might indicate that pair housing improves tokay gecko welfare and suggest that the presence of a conspecific should be considered to improve welfare policies in social reptiles. This study serves as a baseline for future research into how enclosure furnishings, husbandry techniques, and enrichment practices impact the welfare of tokay geckos which will be crucial for refining our understanding of and improving on the welfare of reptiles in captivity.

## Introduction

Research into animal welfare is biased towards mammals, even though a wide range of animals are held in captivity (Doody, 2023). Research into the welfare of captive reptiles and amphibians is particularly underrepresented (Alligood and Leighty, 2015) even though millions of individuals are held in captivity. For example, in 2016, it was estimated that 0.9 million reptiles were kept as pets in the UK alone (PFMA, 2016) which has increased to 1.5 million (PFMA, 2022) and numbers are still increasing worldwide (Moszuti et al., 2017). A lack of information about a reptiles natural history and little research into the welfare consequences of different captive conditions lead to increased mortality within the first year in captivity (Robinson et al., 2015). Therefore, a better understanding of species specific requirements in captivity is urgently needed to mitigate this mortality.

The abiotic environment is an important factor influencing welfare of captive animals (Shepherdson, 1998), possibly more so for ectotherms than endotherms. Contrary to endotherms, like mammals and birds, ectotherms rely on environmental temperature to warm up their body to achieve optimal biological function (Burman et al., 2016; Gillingham and Clark, 2023; Lillywhite, 2023). Thermal preferences can vary greatly across species (Divers and Mader, 2005) highlighting the need for species specific information. Other features of the physical environment such as humidity, availability of shelters, cover or water bodies and climbing opportunities can be equally important (Burman et al., 2016; Divers and Mader, 2005; Lillywhite, 2023). The addition of a range of objects and even olfactory cues has been successful in eliciting desired behaviour or reducing undesired behaviour in reptiles (Divers and Mader, 2005). For example, leopard geckos (*Eublepharis macularius*) show a higher diversity of behaviour in response to thermal, feeding, olfactory, and object enrichments but not in response to visual enrichment (Bashaw et al 2016). However, the welfare of eastern fence lizards (*Sceloporus undulatus*) was not influences by the addition of raised platforms (Rosier and Langkilde, 2011). Therefore, investigating the effect of a range of enrichment methods under different conditions can provide critical information on enclosure design and the possible relationship of different features of the physical environment and welfare (Shepherdson, 1998).

Enrichment might also target the social environment (Shepherdson, 1998), but how the social environment impacts welfare in reptiles is poorly understood (e.g. Tetzlaff et al 2022). One possible explanation for the lack of research in this area is that reptiles have long been viewed as instinct driven (Burghardt, 2013; Font et al. 2023) and asocial creatures that only come together to reproduce (Doody et al. 2013; 2021); a common misconception. A plethora of research has already uncovered the secret social lives of many reptiles species (Doody et al. 2021) and research has shown that even seemingly solitary reptiles are able to learn socially (e.g. Damas-Moreira et al. 2018; Noble et al. 2014; Wilkinson et al 2010) indicating that we still understand too little about what “being social” means in reptiles. We should, therefore, assume that the social environment, especially in species that form stable pairs or live in family groups, is as important as in other social non-reptile species (e.g. Hurst et al. 1997, 1998; Meehan et al. 2003; Williams et al. 2017; Visser et al. 2008). Consequently, research into how social housing influence reptile welfare in captivity is urgently needed.

The aim of this study was to measure changes in the use of enclosure space, basking behaviour and hiding from single to pair housing in captive bred tokay geckos (*Gekko gecko*). Despite Tokay geckos being a popular exotic pet species, only one comprehensive publication exists that summarises information on natural history, behaviour, housing, breeding and husbandry (Grossmann, 2006). It is recommended to keep this species in high enclosures with plenty of hiding spaces and cover, a heat mat rather than a basking spot and in pairs rather than alone (Grossmann, 2006). To date, no empirical work has investigated how tokay geckos use their enclosure space under the recommended conditions and how it is impacted by social housing. Consequently, I empirically investigated how pair housing impacted (1) the frequency with which individuals were found in all areas of their enclosure across time, (2) how basking and hiding were distributed across time, and (3) if males and females differ in these behaviour during single as well as pair housing.

## Methods

### Animals, captive conditions and husbandry

In this study we collected data from 22 captive bred, adult, tokay geckos (*Gekko gecko*, 10 males with Snout Vent Length (SVL) between 12.7-15.5 cm and 12 females with SVL between 11.7-13.9 cm). Sex was determined by the presence (males) or absence (females) of femoral pores (Grossmann, 2006). At the time of the study, lizards were approximately 2-6 years of age and originated from different breeders.

At our facility, individuals are kept in terraria (45 L x 45 B x 70 H cm; 90 L x 45 B x 100 H cm) made out of rigid foam with glass front sliding doors and a mesh top (Figure 1). Enclosures are equipped with a compressed cork back wall, cork branches, refuges made out of cork branches cut in half and hung on the back wall and life plants. We provide UVB light (Exo Terra Reptile UVB 100, 25 W; Figure 1) and a heat mat (TropicShop) fixed on the right outside side wall for thermoregulation (increasing the temperature by 5-10 °C). The ground of the enclosures consists of two layers. The bottom drainage layer is made of expanded clay; the top layer consists of organic rainforest soil (Dragon BIO-Ground). The layers are separated by a mosquito mesh to prevent mixing. Our enclosures are bio-active with collembola, isopods and earth worms in the soil that break down the faecal matter produced by the geckos. We also provide autoclaved red oak leaves and sphagnum moss on top of the soil for the invertebrates to hide in.

**Figure 1.**
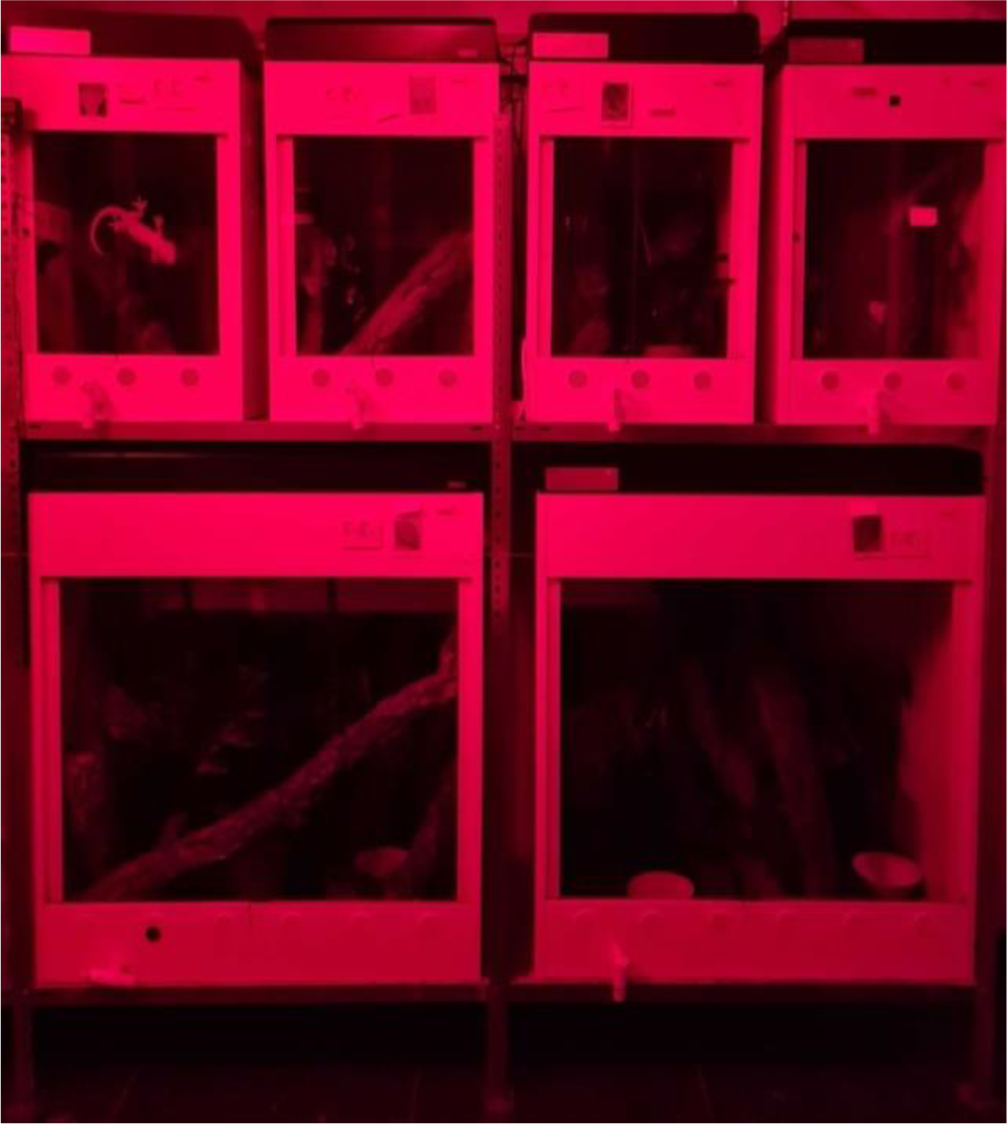
Picture of the housing conditions under red light. Enclosures are made of rigid foam with glass front sliding doors and a mesh top. Small enclosures (45L x 45W x 70H cm) on the top of the shelf housed females during single housing. Large enclosures (90L x 45W x 100H cm) on the bottom of the shelf housed single males or pairs during pair housing. Enclosures were equipped with a cork back wall, cork refuges hung on the back wall with hooks, cork branches, life plants and a water bowl. We also provide a heat mat on the outside of the enclosure and a UVB light source on top.

Enclosures are kept in a fully climate controlled environment that mimics the tropical conditions lizards experience in nature. Room humidity is kept at 50% and daily rainfall (reverse osmosis water, 30s every 12h at 5pm and 4am) increases the humidity within enclosures to 100% for a short period of time. Temperature fluctuates according to the day-night cycle. During the day, enclosure temperature reached 31°C while during the night cycle it reduces to approximately 25 °C. The change in temperature is accompanied by a simulated sunrise and sunset. Temperature and humidity are continuously monitored inside and outside enclosures by the climate system. We keep lizards across two rooms and enclosures are set up on shelves. Small enclosures are kept on top of the shelves and large enclosures on the bottom (Figure 1). Tokay geckos are nocturnal. To work with them during their natural activity period we keep lizards under a reversed 12h:12h photo period (light: 6pm to 6am, dark: 6am to 6pm). A red light (PHILIPS TL-D 36W/15 RED) not visible to geckos (Loew 1994) enables researcher to work with the animals (Figure 1).

Lizards are fed 3-5 adult house crickets (*Acheta domesticus*) three times per week on Monday, Wednesday and Friday. Crickets are gut loaded with cricket mix (reptile planet LDT), Purina Beyond Nature’s Protein^TM^ Adult dry cat food and fresh carrots to ensure that they provided optimal nutrition (Vitamin D and calcium). We feed lizards with 25 cm long forceps to monitor their food intake and use a dim white light (LED, SPYLUX^®^ LEDVANCE 3000K, 0.3 W, 17 lm) to which lizards are conditioned. Water is provided *ad libitum* in water bowls. To keep track of our lizards’ health and general body condition, we weigh them once a month and measure their snout vent length (SVL) every two to three months.

### Experimental setup

I performed two experiments collecting data on the use of enclosure space, basking behaviour and hiding using a within-individual design. Experiment 1 (single housing) was conducted from the 22^nd^ of September to the 3^rd^ of December 2021. During this time, lizards were kept in isolation. All females (N = 12), but one, were kept in small enclosures (45 L x 45 B x 70 H cm; equals more than two – three times SVL) while all males (N = 10), but one, were kept in large enclosures (90 L x 45 B x 100 H cm; equals more than five times SVL). In January 2022, I introduced females to males for the purpose of breeding and established 9 breeding pairs (N = 18 individuals) all kept in large enclosures (90 L x 45 B x 100 H cm). Experiment 2 (pair housing) was conducted from 25^th^ of January to the 28^th^ of April giving females three weeks to adjust to the new housing conditions before data collection.

### Experimental procedure

#### Space use

To measure how lizards use the space within their enclosures during their active period, I recorded their spatial location using scan sampling. On two days per week (Monday to Friday), I entered the animal rooms every 15 minutes for a total of 12 samples. Only one room was sampled at a time and lizards’ behaviour was sampled in a predetermined random order each sampling event. On a given sampling day, I sampled a room either only in the morning between 8:00 and 10:45 or only in the early afternoon between 11:30 and 14:15 (two sampling periods per day). Sampling both rooms at the same time was not feasible due to time constraints. Sampling periods were evenly distributed across rooms and across week days. On feeding days (Monday, Wednesday and Friday) one room was always sampled before feeding while the second room was samples after feeding. Both rooms were sampled on an equal number of feeding and non-feeding days. Overall, I took 12 samples per room per day, two days a week for a total of 10 weeks during single housing and 12 weeks during pair housing resulting in 240/288 data points per individual, respectively (or 5,280/5,616 data points for the whole group, respectively).

During a given sampling event, I entered the room quietly, always from the same door. If possible, a lizards’ location was recorded without approaching an enclosure or using a dim white light. If lizards could not be located from a distance (1.5 meters), its enclosure was approached quietly. If the lizards could still not be located due to the light conditions the white light was briefly turned on and the enclosure searched. If a lizard could still not be found, I opened the enclosure door and used a mirror to check behind big branches and refuges. The duration the white light was used was kept to a minimum to not disturb lizards’ natural behaviour. Each sampling event (whole room) lasted for approximately 2 minutes.

### Data collection

To record the spatial location of an individual during sampling, I divided the whole enclosure into 8 sections: (1) top, left, front; (2) top, right, front; (3) bottom, left, front; (4) bottom, right, front; (5) top, left, back; (6) top, right, back; (7) bottom, left, back; (8) bottom, right, back. These locations were recorded on a coordinate system with left and right on the x-axis, top and bottom on the y-axis and front and back on the z-axis. Left was recorded as 1, and right as 2; top was recorded as 1 and bottom as 2 and front was recorded as 1 and back as 2. Additionally, I recorded if an individual was found under a refuge (cork branches cut in half hanging on the back wall). Starting from week three, I also recorded if a lizard was found on the ceiling, on the ground or on the heat mat (located on the top, right, back side wall). These variables were recorded as presence (1) or absence (0) and were mutually exclusive. During pair housing, I recorded two additional variables related to breeding: (1) if eggs were present in the enclosure (yes = 1 and no = 0) and (2) if an individual was found close to the eggs (within half a body length; yes = 1 and no = 0). Tokay geckos attach their eggs to vertical surfaces such as the back wall behind shelters or branches. Enclosure temperature was automatically recorded every 15 minutes.

### Ethical note

Our scan samples of animal behaviour were strictly non-invasive and followed the guidelines provided by the Association for the Study of Animal Behaviour/ Animal Behaviour Society for the treatment of animals in behavioural research and Teaching (2022), the Guidelines for the ethical use of animals in applied animal behaviour research by the International Society for Applied Ethology (Sherwin et al. 2003) and the International Guiding Principles for Biomedical Research Involving Animals as issued by the Council for the International Organizations of Medical Sciences (2012). Experiments were approved by the Suisse Federal Food Safety and Veterinary Office (National No. 33232, Cantonal No. BE144/2020). Captive conditions were approved by the Suisse Federal Food Safety and Veterinary Office (Laboratory animal husbandry license: No. BE4/11).

### Statistical analyses

All statistical analyses were run in R version 4.2.2 (R Core Team, 2022). I assumed statistical significance if the 95% confidence interval did not cross 0. Data generated during this study and the analysis code are available for download from the Open Science Framework (OSF, link for review purposes: https://osf.io/6v3an/?view_only=0fbe377b4e5e4f7992e6797c25ef9481). No direct comparison of behaviour across single and pair housing was done statistically because only females were moved into new enclosures, and therefore, I was not able to differentiate changes based on being in a novel environment from having a social partner in females.

#### Single housing

To understand geckos spatial behaviour when housed singly, I analysed (1) the probability of moving from one area to another (Bernoulli variable, 1 = moved to another area, 0 = stayed in the same area), (2) the probability to be found hiding behind a shelter (Bernoulli variable, 1 = hiding, 0 = not hiding), (3) the probability to be in the front half of the enclosure (Bernoulli variable, 1 = at the front, 0 = in the back) and (4) the probability to bask on the heat mat (Bernoulli variable, 1 = in the area heated by the mat, 0 = in another area of the enclosure). I used Bayesian generalised linear mixed models (GLMM, one for each variable) using the package *brms* (Brückner 2017; 2018; 2021) with Bernoulli family and logit link function.

As I was interested in differences in behaviour related to testing day, time of day and sex, I included the main effects day (feeding or non-feeding day), time (morning or afternoon) and sex (male or female) and their three-way interaction as fixed effects. Furthermore, as temperature affects behaviour in ectotherms, I also included the main effect of enclosure temperature as a fixed effect. All models included a random intercept of animal identity and a random slope of trial nested in session (i.e. test day) to account for repeated measures. I ensured that model Rhat was 1, that the ESS was above 2000 and checked the density plots and correlation plots to ensure that the models had sampled appropriately. I used a diffuse normal prior with a mean of 0 and a standard deviation of 1 and I ran 4 chains per model of 5000 iterations each. I investigated the results of interactions using *post hoc* least square means comparisons (EMM) from the package *emmeans* (Lenth 2023).

#### Pair housing

To understand geckos spatial behaviour when housed in breeding pairs, I analysed (1) the probability of moving from one area to another (Bernoulli variable, 1 = moved to another area, 0 = stayed in the same area), (2) the probability to be found hiding behind a shelter (Bernoulli variable, 1 = hiding, 0 = not hiding), (3) the probability to be in the front half of the enclosure (Bernoulli variable, 1 = at the front, 0 = in the back), (4) the probability to bask on the heat mat (Bernoulli variable, 1 = in the area heated by the mat, 0 = in another area of the enclosure) and (5) the probability to be close to eggs (Bernoulli variable, 1 = within half a body length of eggs, 0 = outside half a body length of eggs). I used Bayesian generalised linear mixed models (one for each variable) with Bernoulli family and logit link function.

Again, I was interested in differences in behaviour related to testing day, time of day, sex and temperature. Accordingly, all models included the main effects day, time, sex, and temperature as well as the interaction between day and time as fixed effects. Additionally, in the model looking at shelter and heat mat usage, I also included the main effect of egg presence (yes or no) and the interaction between sex and the presence of eggs in the enclosure as a fixed effect. This was done to investigate if the presence of eggs was related to sex specific changes in these two behaviours. All models included a random intercept of animal identity and a random slope of trial nested in session (i.e. test day) to account for repeated measures. I ensured that model Rhat was 1, that the ESS was above 2000 and checked the density plots and correlation plots to ensure that the models had sampled appropriately. I used a diffuse normal prior with a mean of 0 and a standard deviation of 1 and I ran 4 chains per model of 5000 iterations. I investigated the results of interactions using *post hoc* least square means comparisons from the package *emmeans*.

#### Repeatability of behaviour

Apart from changes in behaviour I was also interested in finding out if the behaviours I recorded were repeatable (i.e. consistent within individuals but different across individuals). Consistency in behaviour such as hiding, moving or basking could indicate differences in anxiety or thermal needs which can give important information on enclosure design that takes individuality into account. To this end, I calculated adjusted repeatability using the package rptR (Stoffel et al. 2017) for all behaviours except the probability of being close to the eggs. I ran binary models adjusting for significant predictors (as per the results of the main analysis; day, time, temperature, sex, or eggs present) where appropriate and included a random effect of animal identity.

## Results

### Use of space during single and pair housing

During single housing, individuals spent, on average, 12.5% (range 5.6% - 24.7%) of sampling points in the different sections of their enclosure; this was equal across sexes (Table 1). Similarly, during pair housing, lizards spent, on average, 12.5% (range 2.7% - 28.4%) of sampling points in the different sections of their enclosure. Contrary to single housing, however, males spent more than twice as many sampling points in the top, front of the enclosure than females (Table 1) while females spent almost twice as many sampling points in the top, back part of the enclosure than males (Table 1). Moreover, both during single and pair housing, lizards only spent 0.33% / 0.42% and 1.69% / 0.99% (single / pair housing) of sampling points on the ground or ceiling, respectively (electronic supplementary Table S1).

**Table 1.** Percent of sampling events lizards spent in the eight sections of their enclosure. Data a split into single and pair housing as well as males and females.

|  |  | Single housing |  |  |  | Pair housing |  |  |  |
| --- | --- | --- | --- | --- | --- | --- | --- | --- | --- |
|  |  | Female |  | Male |  | Female |  | Male |  |
|  |  | Left | Right | Left | Right | Left | Right | Left | Right |
| Front | Top | 18.5% | 20.5% | 19.0% | 24.7% | 12.6% | 14.4% | 28.4% | 22.6% |
|  | Bottom | 10.5% | 9.0% | 5.7% | 5.7% | 5.7% | 4.8% | 5.9% | 8.4% |
| Back | Top | 11.0% | 16.1% | 14.0% | 10.7% | 12.5% | 27.8% | 12.3% | 11.4% |
|  | Bottom | 6.4% | 8.0% | 14.6% | 5.6% | 10.9% | 11.3% | 8.3% | 2.7% |

Both during single and pair housing, I found evidence that lizards were more likely to be found in the front half of the enclosure at higher temperatures (single housing: GLMM, estimate = 0.44, CI_low_ = 0.20, CI_up_ = 0.68; electronic supplementary Table S5; pair housing: GLMM, estimate = 0.23, CI_low_ = 0.02, CI_up_ = 0.44; electronic supplementary Table S9). While females and males did not differ in their probability to be found in the front half of the enclosure during single housing (GLMM, estimate = 0.17, CI_low_ = −0.64, CI_up_ = 0.97; Figure 2; electronic supplementary Table S5), during pair housing, males were more likely found in the front compared to females (GLMM, estimate = 1.07, CI_low_ = 0.19, CI_up_ =1.90; Figure 2; electronic supplementary Table S9). *Post hoc* analysis revealed that, during single housing, lizards were more likely found at the front on non-feeding days both in the morning (EMM, estimate = - 0.26, CI_low_ = −0.45, CI_up_ = −0.09) and afternoon (EMM, estimate = −0.41, CI_low_ = −0.59, CI_up_ = - 0.23). However, during pair housing, I found no evidence that lizards probability to be at the front changed based on sampling day and time (morning: EMM, estimate = 0.0.03, CI_low_ = - 0.17, CI_up_ = 0.20; afternoon: EMM, estimate = −0.17, CI_low_ = −0.35, CI_up_ = 0.01).

**Figure 2.**
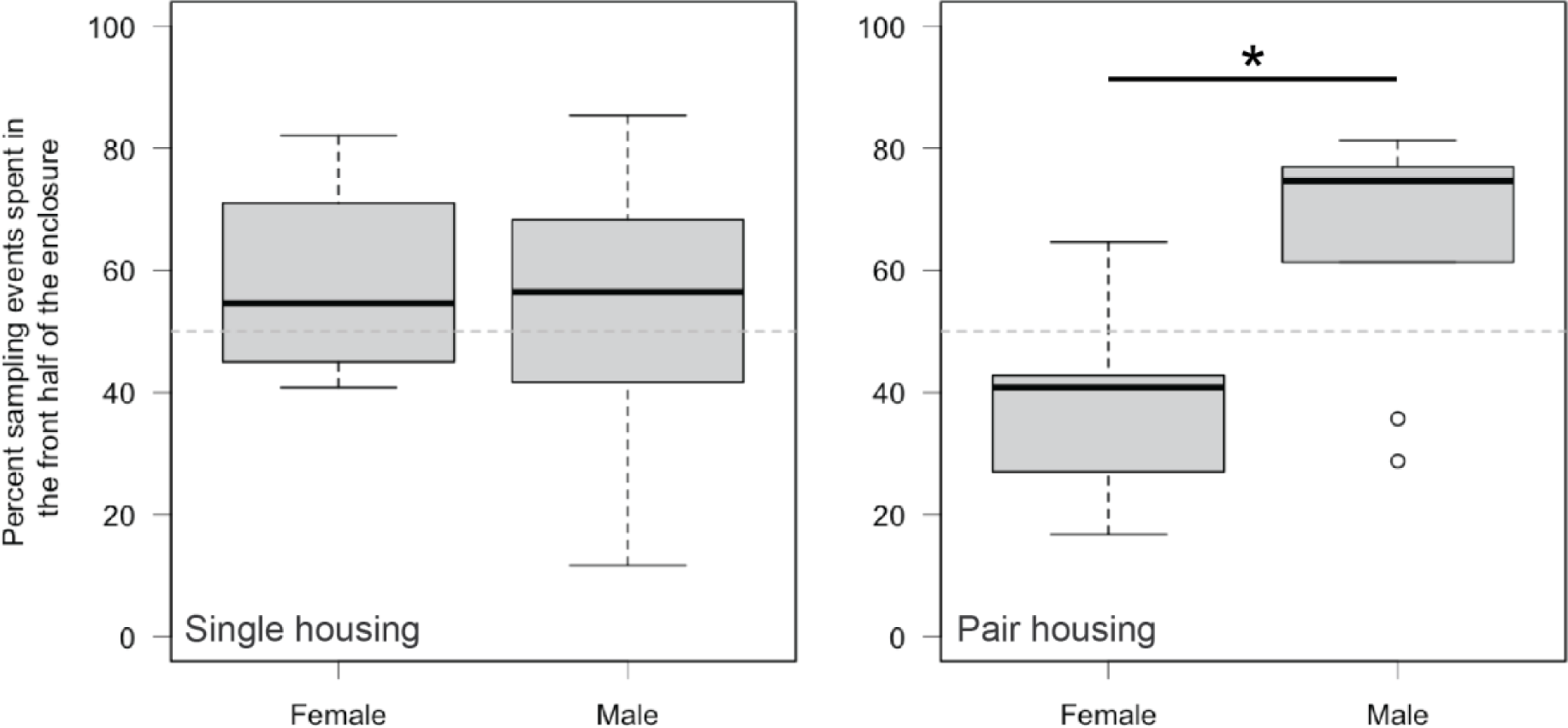
Percent sampling points spent in the front half of the enclosure split by males and females during single and pair housing. * significant difference (confidence interval not crossing 0).

### Probability to move during single and pair housing

During single housing but not pair housing (GLMM, estimate = 0.13, CI_low_ = −0.04, CI_up_ = 0.31; electronic supplementary Table S7), I found evidence that lizards were more likely to move at higher temperatures (GLMM, estimate = 0.45, CI_low_ = 0.24, CI_up_ = 0.67; electronic supplementary Table S3). Neither during single nor pair housing did females and males differ in their probability to move across feeding and non-feeding days or time of day (single housing: GLMM, estimate = −0.01, CI_low_ = −0.49, CI_up_ = 0.46; electronic supplementary Table S3; pair housing: GLMM, estimate = −0.04, CI_low_ = −0.53, CI_up_ = 0.47; electronic supplementary Table S7). Importantly, only during single housing, were lizards more likely to move in the morning, before feeding (EMM, estimate = 0.17, CI_low_ = 0.01, CI_up_ = 0.35; electronic supplementary Figure S1), and less likely to move in the afternoon, after feeding (EMM, estimate = −0.39, CI_low_ = −0.57, CI_up_ = −0.22; electronic supplementary Figure S1). I did not find this pattern during pair housing (GLMM, estimate = −0.15, CI_low_ = −0.40, CI_up_ = 0.09; electronic supplementary Figure S1;) which was confirmed by a *post hoc* test (morning: EMM, estimate = 0.12, CI_low_ = - 0.06, CI_up_ = 0.30; afternoon: EMM, estimate = −0.04, CI_low_ = −0.21, CI_up_ = 0.14).

### Hiding behaviour during single and pair housing

During single housing, I found evidence that lizards were less likely to hide at higher temperatures (GLMM, estimate = −0.88, CI_low_ = −1.24, CI_up_ = −0.53; electronic supplementary Table S4). I also found evidence that males and females differed in the probability to hide across feeding and non-feeding days as well as time of day (Figure 2; electronic supplementary Table S4). Both females and males were more likely to be found hiding on feeding days compared to non-feeding days (females: EMM, estimate = 0.55, CI_low_ = 0.28, CI_up_ = 0.85; males: EMM, estimate = 0.24, CI_low_ = 0.01, CI_up_ = 0.46; Figure 3), whereas only females were less likely to hide in the afternoon (females: EMM, estimate = −0.55, CI_low_ = - 0.84, CI_up_ = −0.24; males: EMM, estimate = 0.18, CI_low_ = −0.07, CI_up_ = 0.43; Figure 3). Contrary, during pair housing, I found no evidence that lizards probability to hide was related to enclosure temperature (GLMM, estimate = 0.10, CI_low_ = −0.18, CI_up_ = 0.38), day and time (GLMM, estimate = −0.22, CI_low_ = −0.57, CI_up_ = 0.13; Figure 3) nor did males and females differ in their hiding behaviour in relation to the presence or absence of eggs in the enclosure (GLMM, estimate = 0.10, CI_low_ = −0.50, CI_up_ = 0.70; electronic supplementary Table S8).

**Figure 3.**
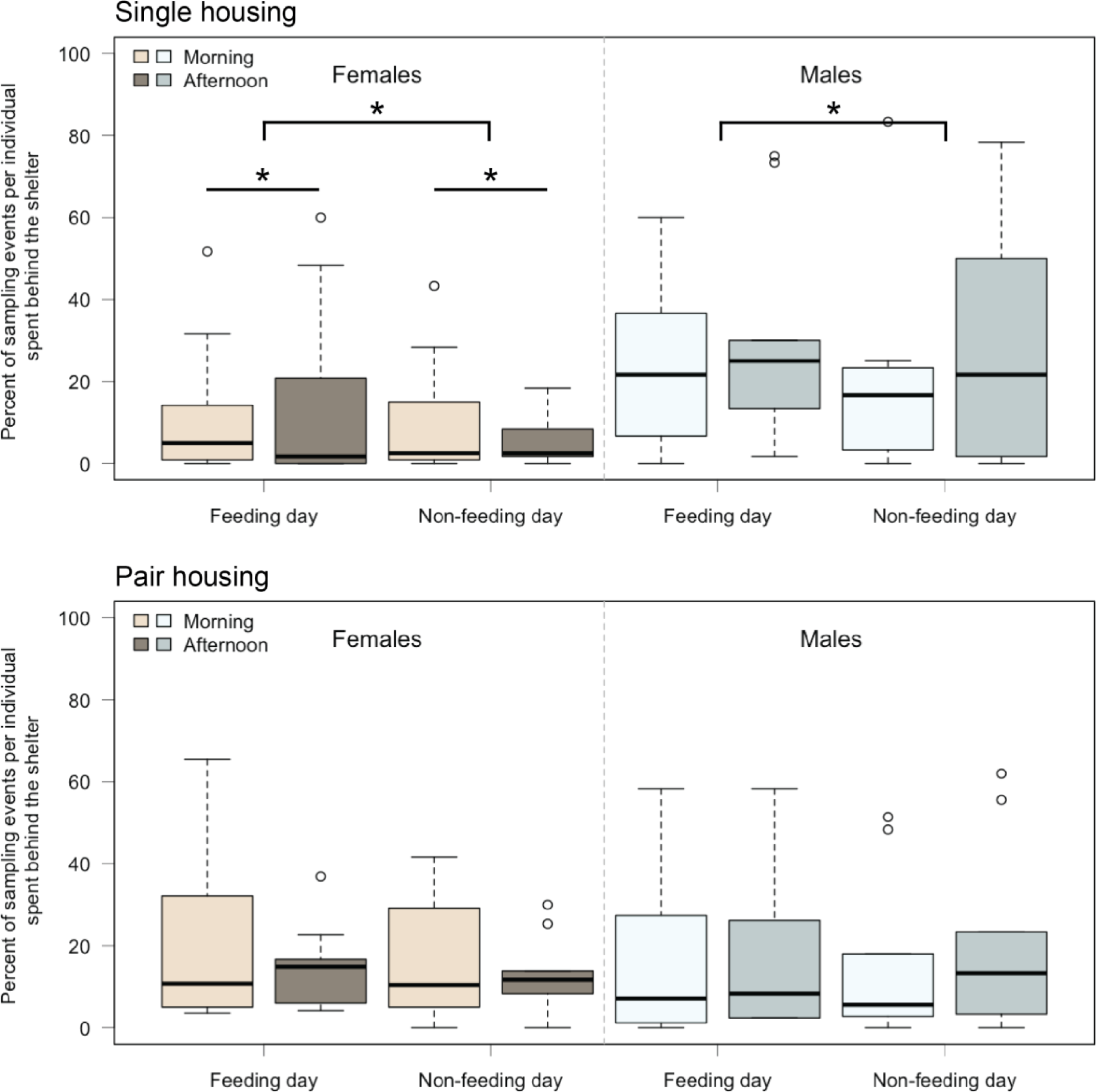
Percent sampling points spent hiding behind a shelter in the morning (light orange and blue) /afternoon (brown and dark blue) on feeding and non-feeding days split by sex (females in orange; males in blue). Top: Behaviour during single housing. Bottom: Behaviour during pair housing. * significant difference (confidence interval not crossing 0). Straight lines indicate difference between morning and afternoon. Hooked lines between feeding and non-feeding days.

### Basking behaviour during single and pair housing

Both during single and pair housing, I found evidence that lizards were less likely to bask on the heat mat at higher enclosure temperatures (single housing: GLMM, estimate = −1.43, CI_low_ = −2.06, CI_up_ = −0.80; electronic supplementary Table S6; pair housing: GLMM, estimate = - 0.39, CI_low_ = −0.71, CI_up_ = −0.08; electronic supplementary Table S10). Furthermore, both during single and pair housing, I found evidence that lizards were more likely to bask in the afternoon on feeding days compared to non-feeding days (single housing: EMM, estimate = 0.84, CI_low_ = 0.47, CI_up_ = 1.20; pair housing: EMM, estimate = 0.50, CI_low_ = 0.26, CI_up_ = 0.74; Figure 4), but not in the morning (single housing: EMM, estimate = 0.13, CI_low_ = −0.26, CI_up_ = 0.50; pair housing: EMM, estimate = −0.32, CI_low_ = −0.60, CI_up_ = −0.04; Figure 4). During single housing, females and males did not differ in their probability to bask across feeding and non-feeding days nor time of day (Figure 5; electronic supplementary Table S6).

**Figure 4.**
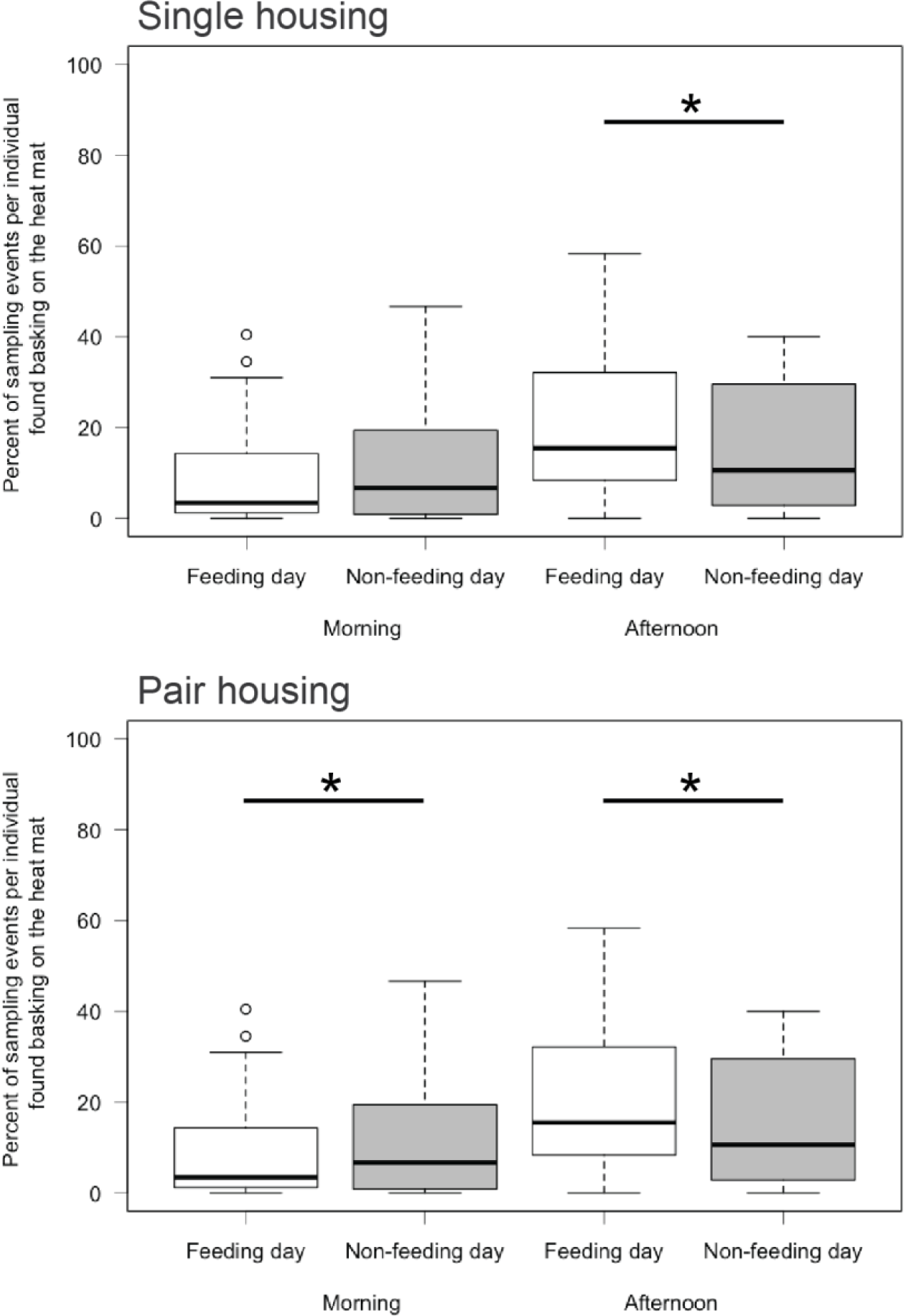
Percent sampling points spent basking on the heat mat in the mourning/ afternoon on feeding (white) and non-feeding days (grey) during single and pair housing. * significant difference (confidence interval not crossing 0).

**Figure 5.**
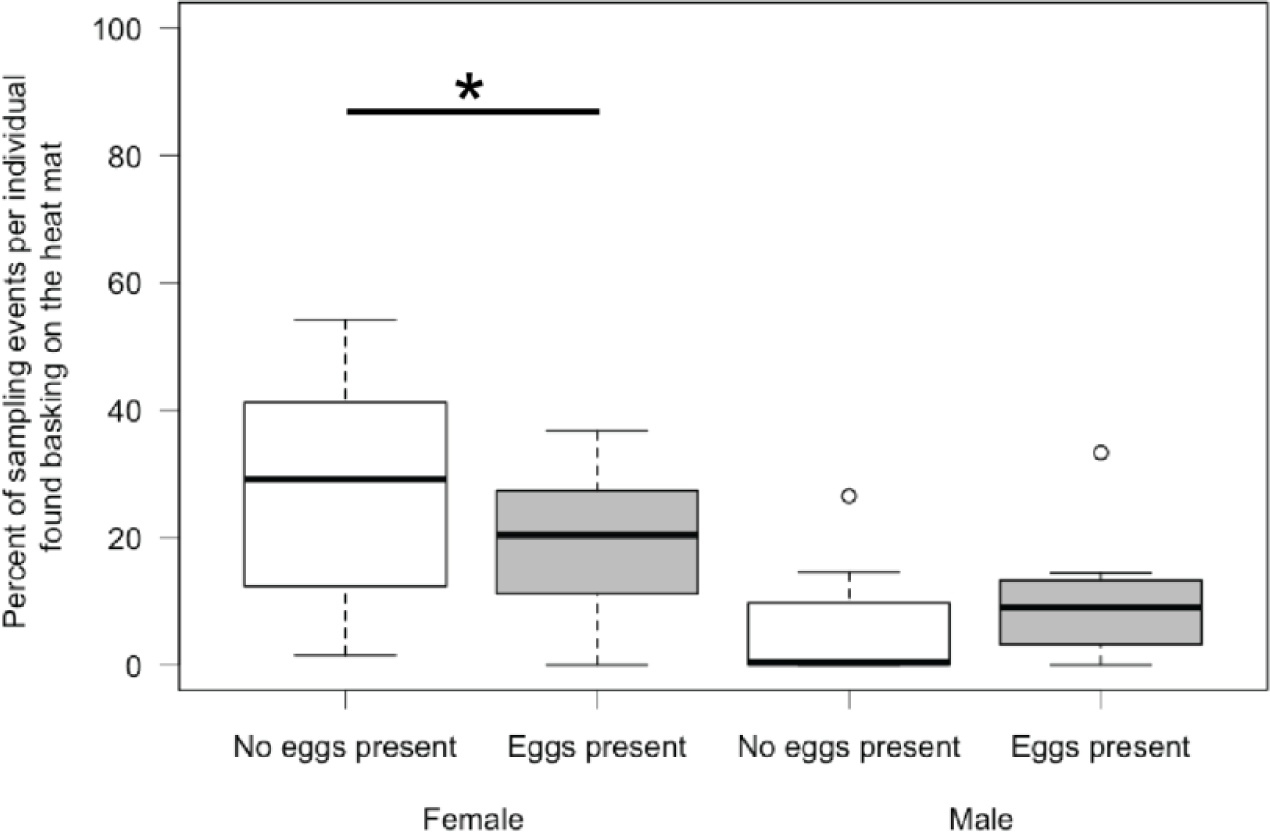
Percent sampling points spent basking on the heat mat of males and females when eggs were not (white) and were present (grey) in the enclosure (during pair housing). * significant difference (confidence interval not crossing 0).

During pair housing, I found no evidence that basking differed across males and females depending on if eggs were present in the enclosure or not (GLMM, estimate = 0.51, CI_low_ = −0.16, CI_up_ = 1.17; Figure 5). However, the main effect of eggs present was significant (GLMM, estimate = −0.74, CI_low_ = −1.16, CI_up_ = −0.33; electronic supplementary Table S10). Consequently, I ran a *post hoc* test that showed that females spent more time basking before their first clutch (EMM, estimate = 0.74, CI_low_ = 0.33, CI_up_ = 1.16; Figure 5) while there was no difference in males (EMM, estimate = 0.23, CI_low_ = −0.35, CI_up_ = 0.79; Figure 5).

### Influence of the presence of eggs in the enclosure

I found no evidence that being found close to eggs was influenced by temperature (GLMM, estimate = −0.17, CI_low_ = −0.69, CI_up_ = 0.37), differed across males and females (GLMM, estimate = −0.49, CI_low_ = −1.97, CI_up_ = 1.05) or day and time (GLMM, estimate = −0.15, CI_low_ = −0.77, CI_up_ = 0.47). However, both main effects of day (GLMM, estimate = 0.61, CI_low_ = 0.09, CI_up_ = 1.12) and time (GLMM, estimate = 0.69, CI_low_ = 0.28, CI_up_ = 1.10; electronic supplementary Table S11) were significant and I ran a *post hoc* test that showed that lizards were more likely found close to eggs on non-feeding days both in the morning (EMM, estimate = −0.45, CI_low_ = −0.89, CI_up_ = −0.01) and afternoon (EMM, estimate = −0.61, CI_low_ = −1.12, CI_up_ = −0.09).

### Repeatability of behaviour

I investigated individual repeatability during single and pair housing in the probability of moving from one area to another, to be found hiding behind a shelter, to be in the front half of the enclosure and to bask on the heat mat. All behaviours were repeatable, but overall, R was below 0.3 (electronic supplementary Table S12). The probability to move was repeatable below 0.1 during both single and pair housing (R_single_ = 0.052; R_pair_ = 0.041) showing high within but low among individual variance. During single housing shelter usage showed the highest repeatability (R = 0.244), while during pair housing heat mat usage showed the highest repeatability (R = 0.289).

## Discussion

Overall, I found changes in enclosure use and basking behaviour form single to pair housing. During single housing, lizards were less likely to be found at the front on feeding days and were also more likely to move in the mornings before feeding compared to the afternoon, behaviour I did not find during pair housing. Furthermore, lizards were more likely to hide on feeding days during single housing but not during pair housing. Overall, environmental temperature affected lizard behaviour but not consistently across the two experiments. While enclosure temperature affected all behaviours measured during single housing which is expected for ectotherms, the probability to move and hide behind a shelter were not related to environmental temperature during pair housing. Consistently, however, lizards were more likely to bask after feeding and females were more likely to bask before their first clutch.

Reptiles are often viewed as asocial animals (Doody et al. 2013; 2021), however, even within lizards, group living has evolved a number of times independently (Halliwell et al. 2017) and the tokay gecko is one of these group living species. In this study, I demonstrate that pair housing has a beneficial impact on tokay geckos similar to species of birds and mammals generally regarded as more social (e.g. Hurst et al. 1997, 1998; Meehan et al. 2003; Williams et al. 2017; Visser et al. 2008). Lizards were more active in the morning and were less likely at the front and hid more on feeding days when housed alone. Predictable husbandry procedures can be related to the emergence of undesirable behaviour such as hyperactivity (Morgan and Tromborg, 2007). I did not find differences in movement and hiding during pair housing, which might indicate a beneficial effect of a mating partner. However, habituation to the captive conditions cannot be ruled out as data collection for the two experiments happened sequentially. Nonetheless, I would expect to find similar increased activity in both experiments because our husbandry procedure did not change. Therefore, it seems likely that the presence of a second individual decreased activity and potentially increased welfare especially considering that hiding behaviour decreased during pair housing. Additionally, I found that shelter usage was more repeatable during single housing than pair housing indicating that individuals were more consistent in their shelter usage during single housing. This result can be interpreted in multiple ways: (1) social facilitation could lead to a decrease in hiding without the second individual influencing fear responses, (2) a mating partner could decrease hiding due to a positive effect on fear responses, or alternatively, (3) instead of hiding, lizards shift to other anti-predator behaviours such as freezing. Further studies on the more detailed influence of a mating partner on fear responses need to be conducted in the future.

Furthermore, males and females differed in their probability to be in the front half of the enclosure during pair housing but not during single housing. For males, the number of sampling points in the front increased while for females it decreased resulting in a difference. This might be related to reproduction. Male tokay geckos are territorial (Grossman, 2006) and, consequently, might have increased the time spent at the front, the transparent part of the enclosure that provided visual access to the room as well as was the only point of access to the enclosure, to protect their territory. However, they show territorial behaviour year round, even when no females are present in their enclosure (personal observation), it is therefore possible that improved welfare due to a mating partner in the enclosure made males more likely to sit at the front. The decrease in females might have been related to egg production and egg deposition on surfaces located in the back of the enclosure. Previous observations in this species from captivity suggested that females are “shyer” because they are less often at the front of the enclosure (Grossman, 2006). The results from this study, together with a previous study showing no difference between males and females in neophobia (Szabo and Ringler, 2023), however, show that the difference between males and females is likely related to reproduction rather than differences in boldness.

My results also demonstrate changes in behaviour in relation to temperature. During pair housing, females were more likely to be found on the heat mat before their first clutch and basking behaviour was more repeatable. Most studies investigating the effect of temperature on embryonal development in lizards focus on the incubation environment after egg deposition (e.g. Oufiero and Angilletta, 2010) and how viviparous females regulate body temperature until parturition (e.g. Uller et al., 2011; Yan et al., 2011). However, temperature also influences embryos within eggs before egg deposition (e.g. Rodríguez-Díaz et al., 2010; Telemeco et al., 2010; Rodríguez-Díaz and Braña, 2011). For example, gravid female *Zootoca vivipara* select lower body temperatures compared to non-gravid females. Incubation of eggs at higher body temperature before deposition leads to more developmentally advances embryos but decreased hatching success and smaller body size and lower running performance after hatching (Rodríguez-Díaz and Braña, 2011). Across studies, females might select lower (e.g. Gainsbury, 2020; Rodríguez-Díaz and Braña, 2011) or higher body temperature (e.g. Juri et al;, 2018; Werner, 1990) when gravid or might not differ from non-gravid females (e.g. Woolrich-Piña et al., 2015). It is possible that increased body temperature is beneficial for egg production and incubation in tokay geckos. Moreover, repeatability could have increased because of females becoming more consistent in heat mat usage due to egg production. However, more controlled studies measuring female body temperature systematically are needed to draw any conclusions about if higher temperatures are important during incubation before oviposition in this species, and if there is a welfare benefit from increased basking opportunity during this time. Furthermore, most of our females were first time breeders and it is possible that the experience of egg production increased stress especially before the first clutch. Short term stress can induce hyperthermia in endotherms (Oka, 2018). To deal with stress, ectotherms need to increase basking to achieve higher body temperature, called behavioural or emotional fever (e.g. Cabanac and Bernieri, 2000; Cabanac and Gosselin, 1993; Preest and Cree, 2008; Rey et al., 2015). If females were actually inducing hyperthermia to deal with stress or if other factors led to increased basking behaviour before deposition of their first clutch has to be investigated in the future.

Finally, lizards were more likely to be found on the heat mat on feeding days in the afternoon. As ectotherms, lizards have to behaviourally thermoregulate to achieve body temperatures that are optimal for biological function including digestion (Burman et al., 2016; Gillingham and Clark, 2023; Lillywhite, 2023). For example, the herbivorous lizard *Cnemidophorus murinus* achieves higher body temperatures for digestion by shifting between microhabitats (Vitt et al., 2005). This behavioural thermoregulation is important because even small changes in body temperature can influence digestion (Troyer, 1987). Together, providing lizards with a constant heat source that is tailored to their needs (e.g. heat spot for diurnal species and a heat mat for nocturnal lizards) is important for their captive welfare to improve digestion as well as potentially help lizards deal with discomfort and stress by allowing them to induce hyperthermia if needed.

## Conclusion

A recent study on the effects of reptile ownership on the perception of reptile cognition and welfare showed that reptile owners had similar low regard for social interactions as a necessary enrichment for captive reptiles as non-owners (Crisante et al., 2023). This impressively demonstrates that even when familiar with reptiles, people still underestimate their social needs. Not surprisingly, the current study is one of the few studies in reptiles (also see Tetzlaff et al., 2022) that took social housing into account when evaluating welfare. Overall, I find decreased movement and hiding behaviour during pair housing that indicate beneficial effects of a mating partner demonstrating the importance of social housing in a social lizard. The data collected in this study function as a baseline from which we can investigate tokay gecko welfare further, to better understand the needs of this species, and consequently, provide them with the best possible captive conditions (Duncan, 1998; Estevez and Christman, 2006).

## Acknowledgements

This work was supported by the Division of Behavioural Ecology at the University of Bern. I want to specifically thank Eva Ringler for her input and support of this study and Océane La Loggia for her insightful comments on an earlier version of this paper.

## Supplementary material

### Supplementary figures

**Figure S1.**
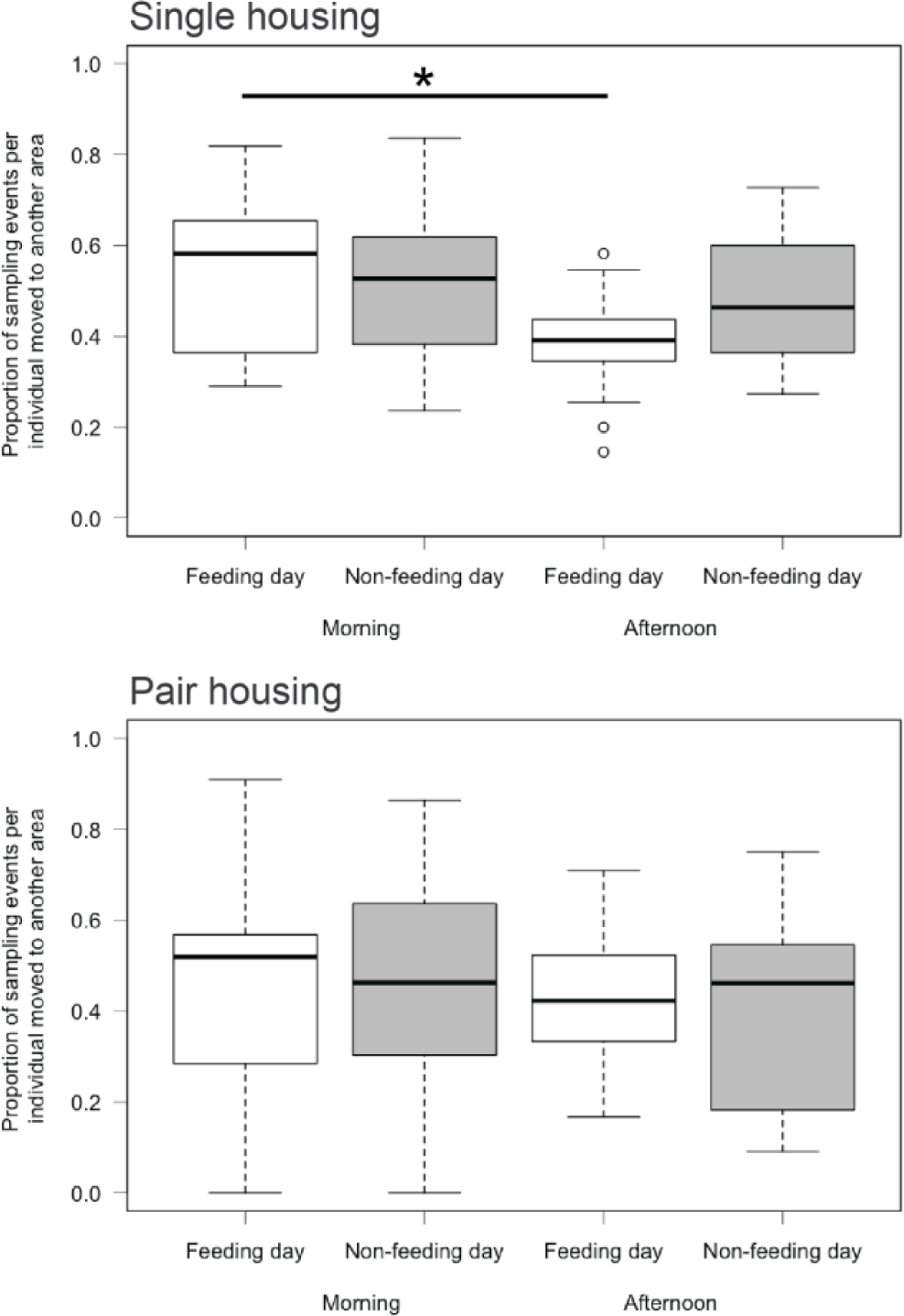
Proportion of sampling points moved from one area of the enclosure to another in the morning/ afternoon on feeding (white) and non-feeding days (grey). Top: Behaviour during single housing. Bottom: Behaviour during pair housing. * significant difference (confidence interval not crossing 0).

### Supplementary tables

**Table S1.**
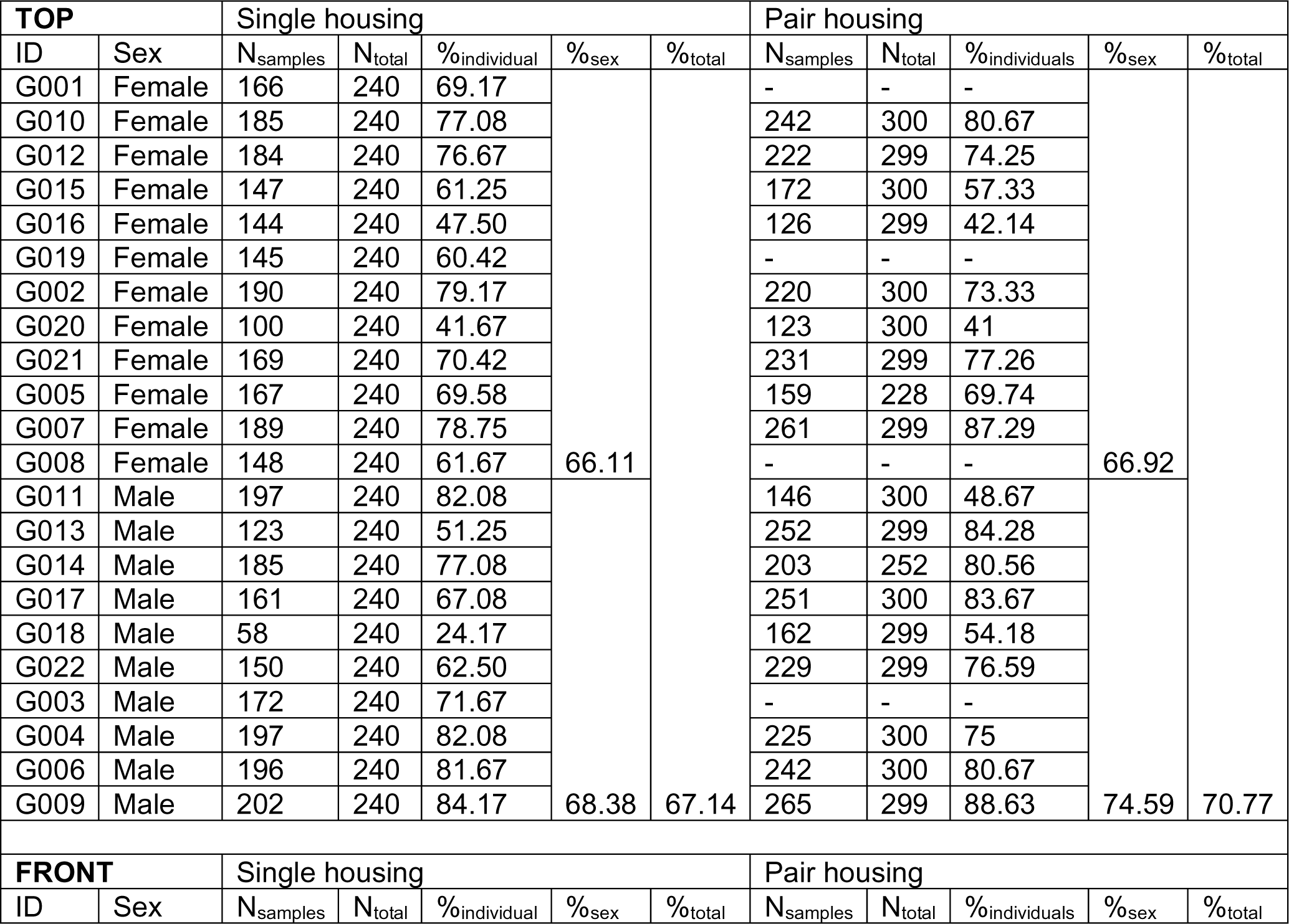
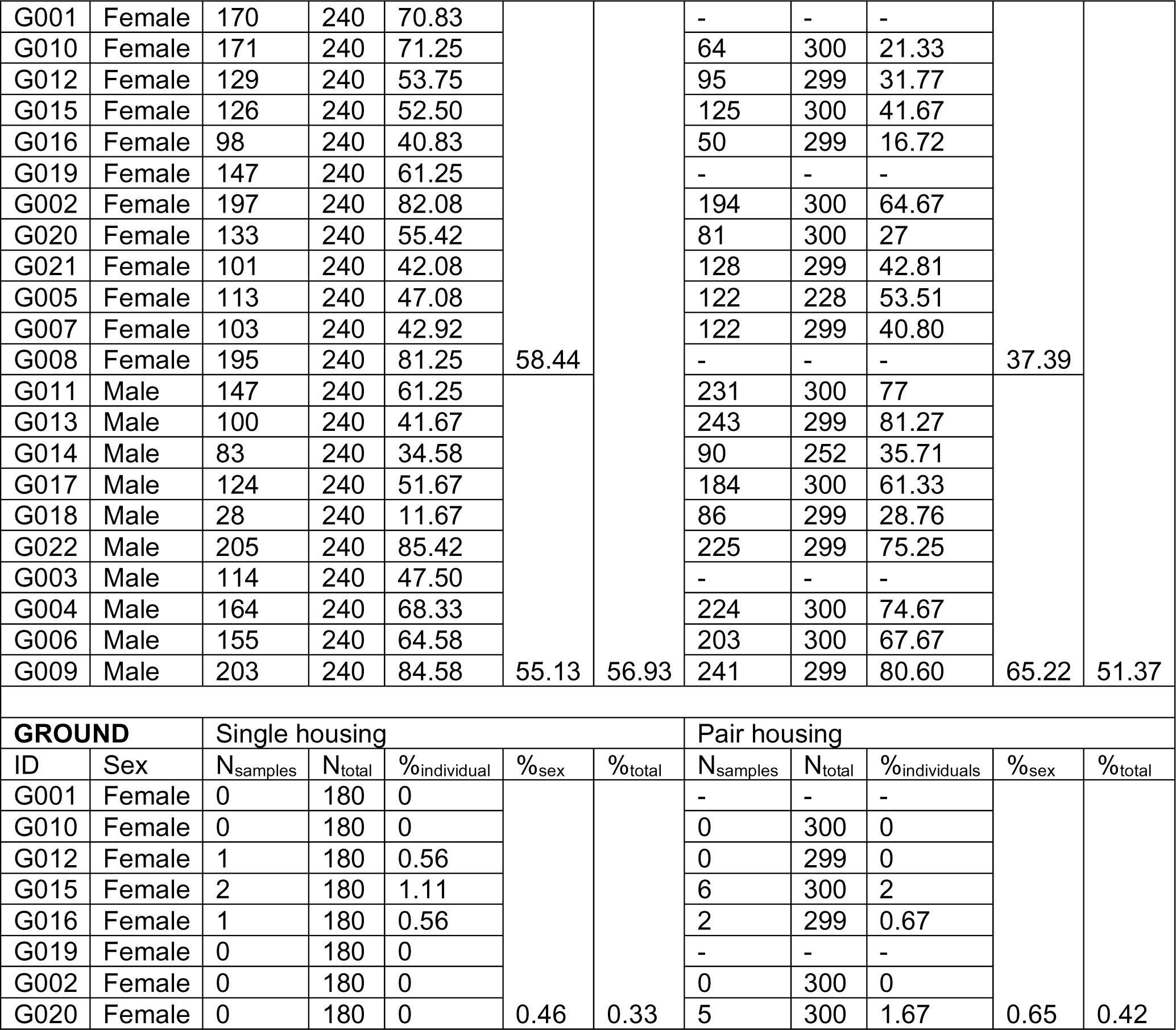
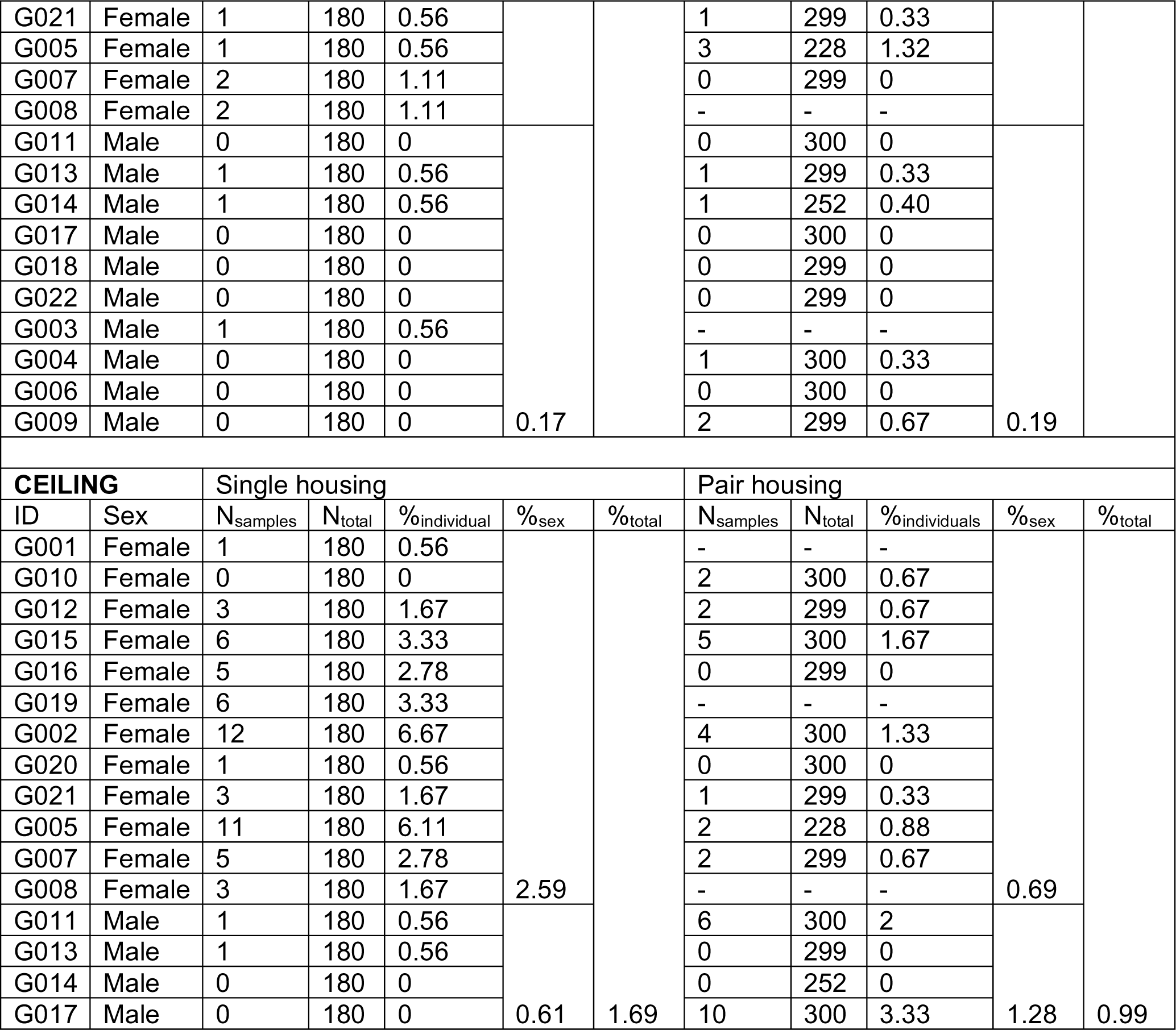
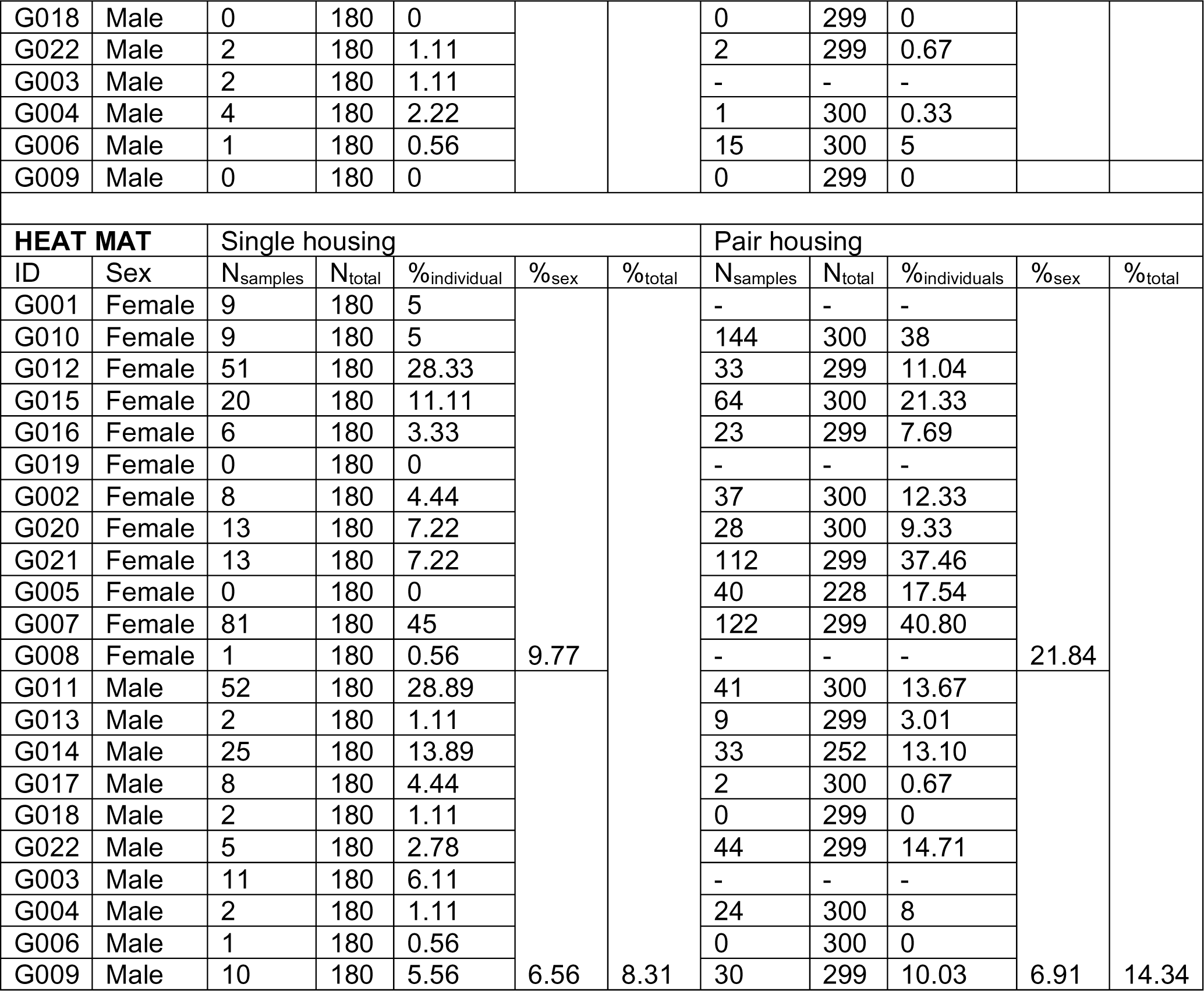
Percent of sampling events individuals were found on the top half, front half, ground or ceiling of their enclosure and basking on the heat mat. ID – individual identity, - no data available (in the second experiment, only those individuals in a pair were observed, N = 18)

| TOP |  | Single housing |  |  |  |  | Pair housing |  |  |  |  |
| --- | --- | --- | --- | --- | --- | --- | --- | --- | --- | --- | --- |
| ID | Sex | N<sub>samples</sub> | N<sub>total</sub> | %individual | %sex | %total | N<sub>samples</sub> | N<sub>total</sub> | %individuals | %sex | %total |
| G001 | Female | 166 | 240 | 69.17 | 66.11 | 67.14 | - | - | - | 66.92 | 70.77 |
| G010 | Female | 185 | 240 | 77.08 |  |  | 242 | 300 | 80.67 |  |  |
| G012 | Female | 184 | 240 | 76.67 |  |  | 222 | 299 | 74.25 |  |  |
| G015 | Female | 147 | 240 | 61.25 |  |  | 172 | 300 | 57.33 |  |  |
| G016 | Female | 144 | 240 | 47.50 |  |  | 126 | 299 | 42.14 |  |  |
| G019 | Female | 145 | 240 | 60.42 |  |  | - | - | - |  |  |
| G002 | Female | 190 | 240 | 79.17 |  |  | 220 | 300 | 73.33 |  |  |
| G020 | Female | 100 | 240 | 41.67 |  |  | 123 | 300 | 41 |  |  |
| G021 | Female | 169 | 240 | 70.42 |  |  | 231 | 299 | 77.26 |  |  |
| G005 | Female | 167 | 240 | 69.58 |  |  | 159 | 228 | 69.74 |  |  |
| G007 | Female | 189 | 240 | 78.75 |  |  | 261 | 299 | 87.29 |  |  |
| G008 | Female | 148 | 240 | 61.67 |  |  | - | - | - |  |  |
| G011 | Male | 197 | 240 | 82.08 | 68.38 | 67.14 | 146 | 300 | 48.67 | 74.59 |  |
| G013 | Male | 123 | 240 | 51.25 |  |  | 252 | 299 | 84.28 |  |  |
| G014 | Male | 185 | 240 | 77.08 |  |  | 203 | 252 | 80.56 |  |  |
| G017 | Male | 161 | 240 | 67.08 |  |  | 251 | 300 | 83.67 |  |  |
| G018 | Male | 58 | 240 | 24.17 |  |  | 162 | 299 | 54.18 |  |  |
| G022 | Male | 150 | 240 | 62.50 |  |  | 229 | 299 | 76.59 |  |  |
| G003 | Male | 172 | 240 | 71.67 |  |  | - | - | - |  |  |
| G004 | Male | 197 | 240 | 82.08 |  |  | 225 | 300 | 75 |  |  |
| G006 | Male | 196 | 240 | 81.67 |  |  | 242 | 300 | 80.67 |  |  |
| G009 | Male | 202 | 240 | 84.17 |  |  | 265 | 299 | 88.63 |  |  |
| FRONT |  | Single housing |  |  |  |  | Pair housing |  |  |  |  |
| ID | Sex | N<sub>samples</sub> | N<sub>total</sub> | %individual | %sex | %total | N<sub>samples</sub> | N<sub>total</sub> | %individuals | %sex | %total |
| G001 | Female | 170 | 240 | 70.83 | 58.44 |  | - | - | - | 37.39 |  |
| G010 | Female | 171 | 240 | 71.25 |  |  | 64 | 300 | 21.33 |  |  |
| G012 | Female | 129 | 240 | 53.75 |  |  | 95 | 299 | 31.77 |  |  |
| G015 | Female | 126 | 240 | 52.50 |  |  | 125 | 300 | 41.67 |  |  |
| G016 | Female | 98 | 240 | 40.83 |  |  | 50 | 299 | 16.72 |  |  |
| G019 | Female | 147 | 240 | 61.25 |  |  | - | - | - |  |  |
| G002 | Female | 197 | 240 | 82.08 |  |  | 194 | 300 | 64.67 |  |  |
| G020 | Female | 133 | 240 | 55.42 |  |  | 81 | 300 | 27 |  |  |
| G021 | Female | 101 | 240 | 42.08 |  |  | 128 | 299 | 42.81 |  |  |
| G005 | Female | 113 | 240 | 47.08 |  |  | 122 | 228 | 53.51 |  |  |
| G007 | Female | 103 | 240 | 42.92 |  |  | 122 | 299 | 40.80 |  |  |
| G008 | Female | 195 | 240 | 81.25 |  |  | - | - | - |  |  |
| G011 | Male | 147 | 240 | 61.25 | 55.13 | 56.93 | 231 | 300 | 77 | 65.22 | 51.37 |
| G013 | Male | 100 | 240 | 41.67 |  |  | 243 | 299 | 81.27 |  |  |
| G014 | Male | 83 | 240 | 34.58 |  |  | 90 | 252 | 35.71 |  |  |
| G017 | Male | 124 | 240 | 51.67 |  |  | 184 | 300 | 61.33 |  |  |
| G018 | Male | 28 | 240 | 11.67 |  |  | 86 | 299 | 28.76 |  |  |
| G022 | Male | 205 | 240 | 85.42 |  |  | 225 | 299 | 75.25 |  |  |
| G003 | Male | 114 | 240 | 47.50 |  |  | - | - | - |  |  |
| G004 | Male | 164 | 240 | 68.33 |  |  | 224 | 300 | 74.67 |  |  |
| G006 | Male | 155 | 240 | 64.58 |  |  | 203 | 300 | 67.67 |  |  |
| G009 | Male | 203 | 240 | 84.58 |  |  | 241 | 299 | 80.60 |  |  |
| GROUND |  | Single housing |  |  |  |  | Pair housing |  |  |  |  |
| ID | Sex | N<sub>samples</sub> | N<sub>total</sub> | %individual | %<sub>sex</sub> | %<sub>total</sub> | N<sub>samples</sub> | N<sub>total</sub> | %individuals | %<sub>sex</sub> | %<sub>total</sub> |
| G001 | Female | 0 | 180 | 0 | 0.46 | 0.33 | - | - | - | 0.65 | 0.42 |
| G010 | Female | 0 | 180 | 0 |  |  | 0 | 300 | 0 |  |  |
| G012 | Female | 1 | 180 | 0.56 |  |  | 0 | 299 | 0 |  |  |
| G015 | Female | 2 | 180 | 1.11 |  |  | 6 | 300 | 2 |  |  |
| G016 | Female | 1 | 180 | 0.56 |  |  | 2 | 299 | 0.67 |  |  |
| G019 | Female | 0 | 180 | 0 |  |  | - | - | - |  |  |
| G002 | Female | 0 | 180 | 0 |  |  | 0 | 300 | 0 |  |  |
| G020 | Female | 0 | 180 | 0 |  |  | 5 | 300 | 1.67 |  |  |
| G021 | Female | 1 | 180 | 0.56 |  |  | 1 | 299 | 0.33 |  |  |
| G005 | Female | 1 | 180 | 0.56 |  |  | 3 | 228 | 1.32 |  |  |
| G007 | Female | 2 | 180 | 1.11 |  |  | 0 | 299 | 0 |  |  |
| G008 | Female | 2 | 180 | 1.11 |  |  | - | - | - |  |  |
| G011 | Male | 0 | 180 | 0 | 0.17 |  | 0 | 300 | 0 | 0.19 |  |
| G013 | Male | 1 | 180 | 0.56 |  |  | 1 | 299 | 0.33 |  |  |
| G014 | Male | 1 | 180 | 0.56 |  |  | 1 | 252 | 0.40 |  |  |
| G017 | Male | 0 | 180 | 0 |  |  | 0 | 300 | 0 |  |  |
| G018 | Male | 0 | 180 | 0 |  |  | 0 | 299 | 0 |  |  |
| G022 | Male | 0 | 180 | 0 |  |  | 0 | 299 | 0 |  |  |
| G003 | Male | 1 | 180 | 0.56 |  |  | - | - | - |  |  |
| G004 | Male | 0 | 180 | 0 |  |  | 1 | 300 | 0.33 |  |  |
| G006 | Male | 0 | 180 | 0 |  |  | 0 | 300 | 0 |  |  |
| G009 | Male | 0 | 180 | 0 |  |  | 2 | 299 | 0.67 |  |  |
| CEILING |  | Single housing |  |  |  |  | Pair housing |  |  |  |  |
| ID | Sex | N<sub>samples</sub> | N<sub>total</sub> | %<sub>individual</sub> | %<sub>sex</sub> | %<sub>total</sub> | N<sub>samples</sub> | N<sub>total</sub> | %<sub>individuals</sub> | %<sub>sex</sub> | %<sub>total</sub> |
| G001 | Female | 1 | 180 | 0.56 | 2.59 | 1.69 | - | - | - | 0.69 | 0.99 |
| G010 | Female | 0 | 180 | 0 |  |  | 2 | 300 | 0.67 |  |  |
| G012 | Female | 3 | 180 | 1.67 |  |  | 2 | 299 | 0.67 |  |  |
| G015 | Female | 6 | 180 | 3.33 |  |  | 5 | 300 | 1.67 |  |  |
| G016 | Female | 5 | 180 | 2.78 |  |  | 0 | 299 | 0 |  |  |
| G019 | Female | 6 | 180 | 3.33 |  |  | - | - | - |  |  |
| G002 | Female | 12 | 180 | 6.67 |  |  | 4 | 300 | 1.33 |  |  |
| G020 | Female | 1 | 180 | 0.56 |  |  | 0 | 300 | 0 |  |  |
| G021 | Female | 3 | 180 | 1.67 |  |  | 1 | 299 | 0.33 |  |  |
| G005 | Female | 11 | 180 | 6.11 |  |  | 2 | 228 | 0.88 |  |  |
| G007 | Female | 5 | 180 | 2.78 |  |  | 2 | 299 | 0.67 |  |  |
| G008 | Female | 3 | 180 | 1.67 |  |  | - | - | - |  |  |
| G011 | Male | 1 | 180 | 0.56 |  |  | 6 | 300 | 2 |  |  |
| G013 | Male | 1 | 180 | 0.56 |  |  | 0 | 299 | 0 |  |  |
| G014 | Male | 0 | 180 | 0 |  |  | 0 | 252 | 0 |  |  |
| G017 | Male | 0 | 180 | 0 | 0.61 | 1.69 | 10 | 300 | 3.33 | 1.28 | 0.99 |
| G018 | Male | 0 | 180 | 0 |  |  | 0 | 299 | 0 |  |  |
| G022 | Male | 2 | 180 | 1.11 |  |  | 2 | 299 | 0.67 |  |  |
| G003 | Male | 2 | 180 | 1.11 |  |  | - | - | - |  |  |
| G004 | Male | 4 | 180 | 2.22 |  |  | 1 | 300 | 0.33 |  |  |
| G006 | Male | 1 | 180 | 0.56 |  |  | 15 | 300 | 5 |  |  |
| G009 | Male | 0 | 180 | 0 |  |  | 0 | 299 | 0 |  |  |
| **HEAT MAT** |  | Single housing |  |  |  |  | Pair housing |  |  |  |  |
| ID | Sex | N<sub>samples</sub> | N<sub>total</sub> | %individual | %sex | %total | N<sub>samples</sub> | N<sub>total</sub> | %individuals | %sex | %total |
| G001 | Female | 9 | 180 | 5 |  |  | - | - | - |  |  |
| G010 | Female | 9 | 180 | 5 |  |  | 144 | 300 | 38 |  |  |
| G012 | Female | 51 | 180 | 28.33 |  |  | 33 | 299 | 11.04 |  |  |
| G015 | Female | 20 | 180 | 11.11 |  |  | 64 | 300 | 21.33 |  |  |
| G016 | Female | 6 | 180 | 3.33 |  |  | 23 | 299 | 7.69 |  |  |
| G019 | Female | 0 | 180 | 0 |  |  | - | - | - |  |  |
| G002 | Female | 8 | 180 | 4.44 |  |  | 37 | 300 | 12.33 |  |  |
| G020 | Female | 13 | 180 | 7.22 |  |  | 28 | 300 | 9.33 |  |  |
| G021 | Female | 13 | 180 | 7.22 |  |  | 112 | 299 | 37.46 |  |  |
| G005 | Female | 0 | 180 | 0 |  |  | 40 | 228 | 17.54 |  |  |
| G007 | Female | 81 | 180 | 45 |  |  | 122 | 299 | 40.80 |  |  |
| G008 | Female | 1 | 180 | 0.56 | 9.77 |  | - | - | - | 21.84 |  |
| G011 | Male | 52 | 180 | 28.89 |  |  | 41 | 300 | 13.67 |  |  |
| G013 | Male | 2 | 180 | 1.11 |  |  | 9 | 299 | 3.01 |  |  |
| G014 | Male | 25 | 180 | 13.89 |  |  | 33 | 252 | 13.10 |  |  |
| G017 | Male | 8 | 180 | 4.44 |  |  | 2 | 300 | 0.67 |  |  |
| G018 | Male | 2 | 180 | 1.11 |  |  | 0 | 299 | 0 |  |  |
| G022 | Male | 5 | 180 | 2.78 |  |  | 44 | 299 | 14.71 |  |  |
| G003 | Male | 11 | 180 | 6.11 |  |  | - | - | - |  |  |
| G004 | Male | 2 | 180 | 1.11 |  |  | 24 | 300 | 8 |  |  |
| G006 | Male | 1 | 180 | 0.56 |  |  | 0 | 300 | 0 |  |  |
| G009 | Male | 10 | 180 | 5.56 | 6.56 | 8.31 | 30 | 299 | 10.03 | 6.91 | 14.34 |

**Table S2.**
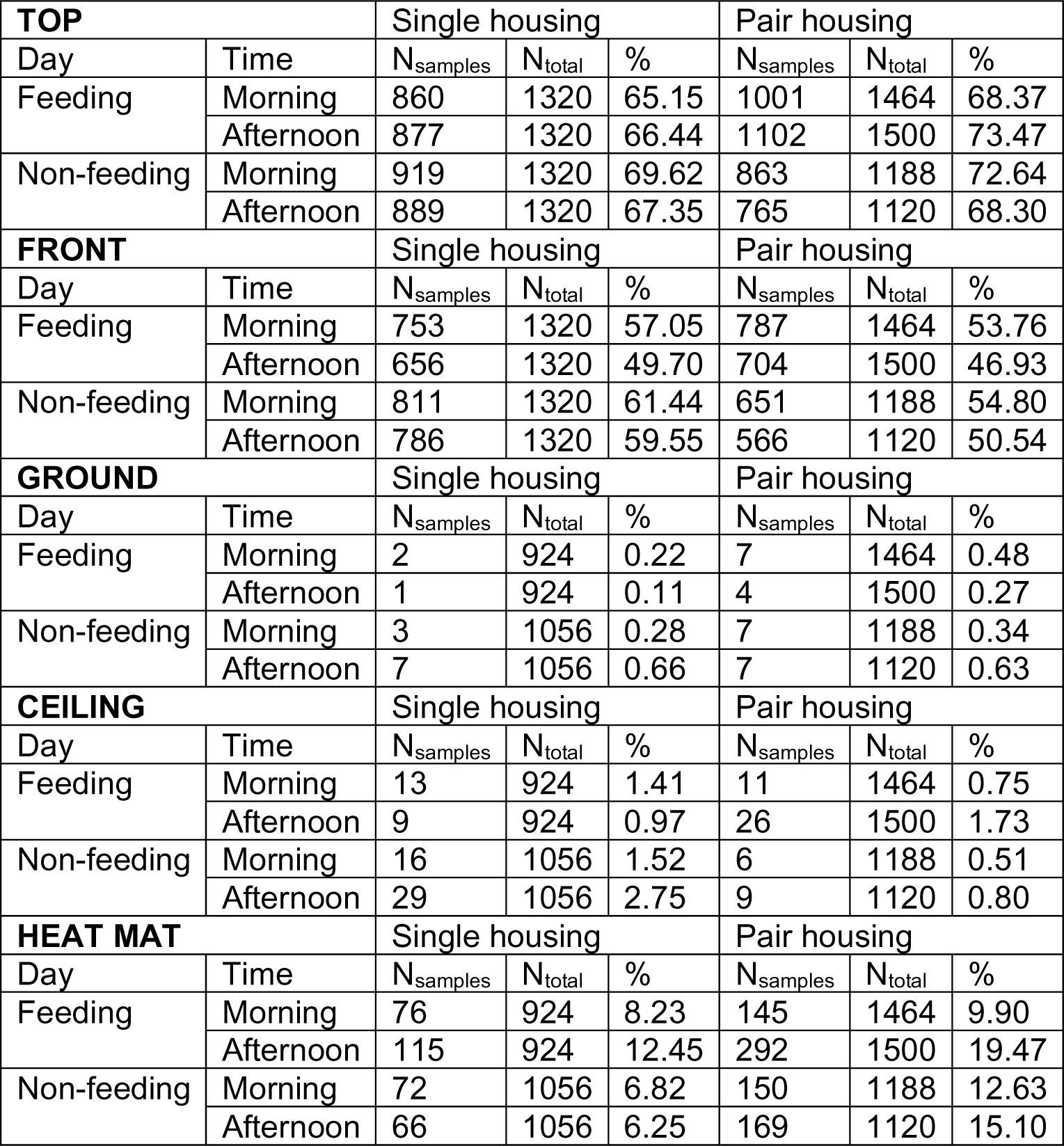
Percent of sampling events individuals were found on the top half, front half, ground or ceiling of their enclosure and basking on the heat mat separated into morning and afternoon on feeding and non-feeding days. ID – individual identity, - no data available (in the second experiment, only those individuals in a pair were observed, N = 18)

| **TOP** |  | Single housing |  |  | Pair housing |  |  |
| --- | --- | --- | --- | --- | --- | --- | --- |
| Day | Time | N<sub>samples</sub> | N<sub>total</sub> | % | N<sub>samples</sub> | N<sub>total</sub> | % |
| Feeding | Morning | 860 | 1320 | 65.15 | 1001 | 1464 | 68.37 |
|  | Afternoon | 877 | 1320 | 66.44 | 1102 | 1500 | 73.47 |
| Non-feeding | Morning | 919 | 1320 | 69.62 | 863 | 1188 | 72.64 |
|  | Afternoon | 889 | 1320 | 67.35 | 765 | 1120 | 68.30 |
| **FRONT** |  | Single housing |  |  | Pair housing |  |  |
| Day | Time | N<sub>samples</sub> | N<sub>total</sub> | % | N<sub>samples</sub> | N<sub>total</sub> | % |
| Feeding | Morning | 753 | 1320 | 57.05 | 787 | 1464 | 53.76 |
|  | Afternoon | 656 | 1320 | 49.70 | 704 | 1500 | 46.93 |
| Non-feeding | Morning | 811 | 1320 | 61.44 | 651 | 1188 | 54.80 |
|  | Afternoon | 786 | 1320 | 59.55 | 566 | 1120 | 50.54 |
| **GROUND** |  | Single housing |  |  | Pair housing |  |  |
| Day | Time | N<sub>samples</sub> | N<sub>total</sub> | % | N<sub>samples</sub> | N<sub>total</sub> | % |
| Feeding | Morning | 2 | 924 | 0.22 | 7 | 1464 | 0.48 |
|  | Afternoon | 1 | 924 | 0.11 | 4 | 1500 | 0.27 |
| Non-feeding | Morning | 3 | 1056 | 0.28 | 7 | 1188 | 0.34 |
|  | Afternoon | 7 | 1056 | 0.66 | 7 | 1120 | 0.63 |
| **CEILING** |  | Single housing |  |  | Pair housing |  |  |
| Day | Time | N<sub>samples</sub> | N<sub>total</sub> | % | N<sub>samples</sub> | N<sub>total</sub> | % |
| Feeding | Morning | 13 | 924 | 1.41 | 11 | 1464 | 0.75 |
|  | Afternoon | 9 | 924 | 0.97 | 26 | 1500 | 1.73 |
| Non-feeding | Morning | 16 | 1056 | 1.52 | 6 | 1188 | 0.51 |
|  | Afternoon | 29 | 1056 | 2.75 | 9 | 1120 | 0.80 |
| **HEAT MAT** |  | Single housing |  |  | Pair housing |  |  |
| Day | Time | N<sub>samples</sub> | N<sub>total</sub> | % | N<sub>samples</sub> | N<sub>total</sub> | % |
| Feeding | Morning | 76 | 924 | 8.23 | 145 | 1464 | 9.90 |
|  | Afternoon | 115 | 924 | 12.45 | 292 | 1500 | 19.47 |
| Non-feeding | Morning | 72 | 1056 | 6.82 | 150 | 1188 | 12.63 |
|  | Afternoon | 66 | 1056 | 6.25 | 169 | 1120 | 15.10 |

**Table S3.** Estimates and test statistics from the Bayesian generalised linear model looking at the change in the probability to move from one enclosure area to another during single housing. Significant results (95% confidence interval – CI – not crossing 0) are highlighted in bold.

| Parameter | Estimate | Upper 95% CI | Lower 95% CI |
| --- | --- | --- | --- |
| **Intercept** | **-11.70** | **-16.91** | **-6.33** |
| **Non-feeding day** | **0.45** | **0.22** | **0.68** |
| **Morning** | **0.66** | **0.41** | **0.92** |
| Male | -0.01 | -0.49 | 0.46 |
| **Temperature** | **0.45** | **0.24** | **0.67** |
| **Interaction: day & time** | **-0.56** | **-0.88** | **-0.24** |
| Interaction: day & sex | -0.12 | -0.47 | 0.22 |
| Interaction: time & sex | -0.28 | -0.64 | 0.07 |
| Interaction: day & time & sex | -0.00 | -0.48 | 0.49 |

**Table S4.** Estimates and test statistics from the Bayesian generalised linear model looking at the change in the probability of hiding behind a shelter during single housing. Significant results (95% confidence interval – CI – not crossing 0) are highlighted in bold.

| Parameter | Estimate | Upper 95% CI | Lower 95% CI |
| --- | --- | --- | --- |
| **Intercept** | **19.13** | **10.56** | **27.86** |
| **Non-feeding day** | **-0.93** | **-1.33** | **-0.52** |
| Morning | 0.17 | -0.20 | 0.55 |
| **Male** | **1.26** | **0.13** | **2.36** |
| **Temperature** | **-0.88** | **-1.24** | **-0.53** |
| **Interaction: day & time** | **0.75** | **0.21** | **1.28** |
| **Interaction: day & sex** | **0.85** | **0.38** | **1.34** |
| Interaction: time & sex | -0.19 | -0.64 | 0.27 |
| **Interaction: day & time & sex** | **-1.07** | **-1.74** | **-0.41** |

**Table S5.** Estimates and test statistics from the Bayesian generalised linear model looking at the change in the probability of being in the front half of the enclosure during single housing. Significant results (95% confidence interval – CI – not crossing 0) are highlighted in bold.

| Parameter | Estimate | Upper 95% CI | Lower 95% CI |
| --- | --- | --- | --- |
| **Intercept** | **-10.79** | **-16.82** | **-4.74** |
| **Non-feeding day** | **0.56** | **0.33** | **0.79** |
| **Morning** | **0.25** | **0.00** | **0.50** |
| Male | 0.17 | -0.64 | 0.97 |
| **Temperature** | **0.44** | **0.20** | **0.68** |
| Interaction: day & time | -0.25 | -0.57 | 0.08 |
| Interaction: day & sex | -0.31 | -0.64 | 0.03 |
| Interaction: time & sex | -0.20 | -0.56 | 0.15 |
| Interaction: day & time & sex | 0.21 | -0.27 | 0.70 |

**Table S6.** Estimates and test statistics from the Bayesian generalised linear model looking at the change in the probability of being found basking on the heat mat during single housing. Significant results (95% confidence interval – CI – not crossing 0) are highlighted in bold.

| Parameter | Estimate | Upper 95% CI | Lower 95% CI |
| --- | --- | --- | --- |
| **Intercept** | **32.88** | **17.63** | **48.61** |
| **Non-feeding day** | **-0.60** | **-1.03** | **-0.19** |
| Morning | -0.27 | -0.71 | 0.16 |
| Male | 0.17 | -1.03 | 1.39 |
| **Temperature** | **-1.43** | **-2.06** | **-0.80** |
| **Interaction: day & time** | **0.72** | **0.16** | **1.30** |
| Interaction: day & sex | -0.49 | -1.15 | 0.15 |
| Interaction: time & sex | 0.25 | -0.38 | 0.87 |
| Interaction: day & time & sex | -0.02 | -0.89 | 0.87 |

**Table S7.**
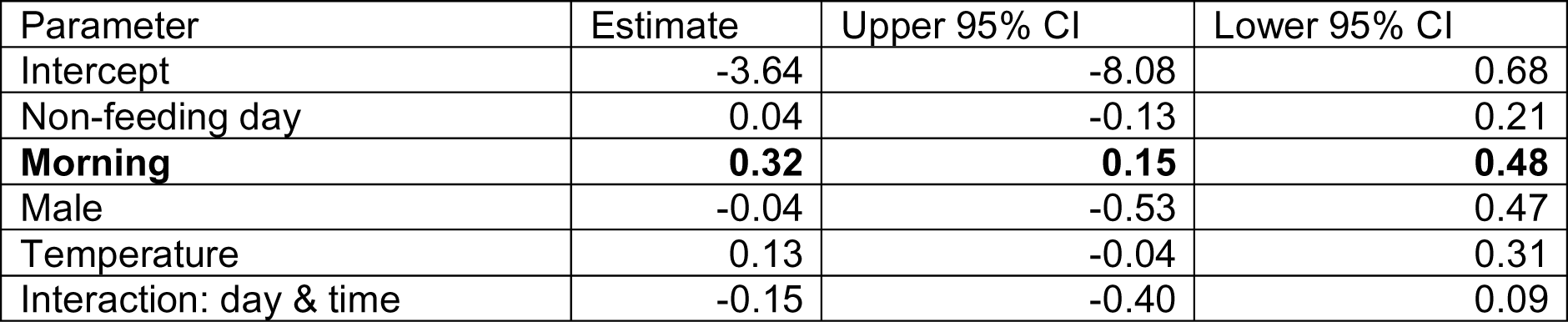
Estimates and test statistics from the Bayesian generalised linear model looking at the change in the probability to move from one enclosure area to another during pair housing. Significant results (95% confidence interval – CI – not crossing 0) are highlighted in bold.

**Table S8.** Estimates and test statistics from the Bayesian generalised linear model looking at the change in the probability of hiding behind a shelter during pair housing. Significant results (95% confidence interval – CI – not crossing 0) are highlighted in bold.

| Parameter | Estimate | Upper 95% CI | Lower 95% CI |
| --- | --- | --- | --- |
| Intercept | -4.42 | -11.38 | 2.46 |
| Non-feeding day | -0.08 | -0.33 | 0.17 |
| Morning | 0.23 | -0.00 | 0.46 |
| Male | -0.42 | -1.53 | 0.69 |
| Eggs present | 0.20 | -0.22 | 0.62 |
| Temperature | 0.10 | -0.18 | 0.38 |
| Interaction: day & time | -0.22 | -0.57 | 0.13 |
| Interaction: sex & eggs present | 0.10 | -0.50 | 0.70 |

**Table S9.**
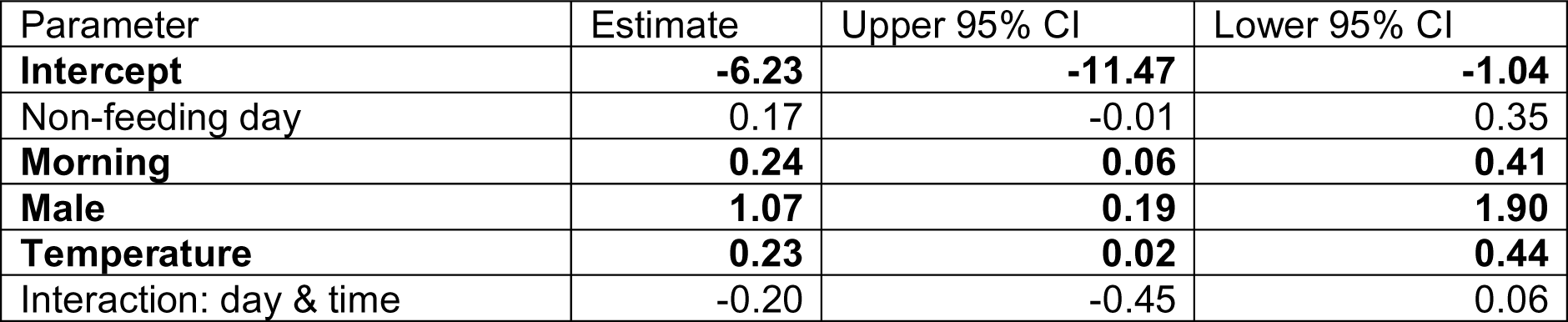
Estimates and test statistics from the Bayesian generalised linear model looking at the change in the probability of being in the front half of the enclosure during pair housing. Significant results (95% confidence interval – CI – not crossing 0) are highlighted in bold.

**Table S10.** Estimates and test statistics from the Bayesian generalised linear model looking at the change in the probability of being found basking on the heat mat during pair housing. Significant results (95% confidence interval – CI – not crossing 0) are highlighted in bold.

| Parameter | Estimate | Upper 95% CI | Lower 95% CI |
| --- | --- | --- | --- |
| **Intercept** | **8.60** | **1.05** | **16.37** |
| **Non-feeding day** | **-0.50** | **-0.74** | **-0.26** |
| **Morning** | **-0.77** | **-1.01** | **-0.52** |
| Male | -1.26 | -2.57 | 0.12 |
| **Eggs present** | **-0.74** | **-1.16** | **-0.33** |
| **Temperature** | **-0.39** | **-0.71** | **-0.08** |
| **Interaction: day & time** | **0.81** | **0.46** | **1.17** |
| Interaction: sex & eggs present | 0.51 | -0.16 | 1.20 |

**Table S11.** Estimates and test statistics from the Bayesian generalised linear model looking at the change in the probability to be found close to eggs during pair housing. Significant results (95% confidence interval – CI – not crossing 0) are highlighted in bold.

| Parameter | Estimate | Upper 95% CI | Lower 95% CI |
| --- | --- | --- | --- |
| Intercept | 2.52 | -10.38 | 15.05 |
| **Non-feeding day** | **0.61** | **0.09** | **1.12** |
| **Morning** | **0.69** | **0.28** | **1.10** |
| Male | -0.49 | -1.97 | 1.05 |
| Temperature | -0.17 | -0.69 | 0.37 |
| Interaction: day & time | -0.15 | -0.77 | 0.47 |

**Table S12.** Model specifics and results from the analysis looking at repeatability in behaviour. R – repeatability, CI_low_ – lower 95% confidence interval, CI_up_ – upper 95% confidence interval. Movement – probability to move between areas in the enclosure, Shelter – probability of hiding behind a shelter, Front – probability to be in the front half of the enclosure, heatmat – probability to be found basking. Significant R are highlighted in bold.

| **Single housing** |  |  |  |
| --- | --- | --- | --- |
| **Model** | **R** | **$CI_{low}$** | **$CI_{up}$** |
| Movement ~ temperature + day + time + (1 \| ID) | **0.052** | 0.022 | 0.083 |
| Shelter ~ sex + temperature + day + (1\|ID) | **0.244** | 0.097 | 0.384 |
| Front ~ temperature + day + time + (1 \| ID) | **0.175** | 0.090 | 0.264 |
| Heatmat ~ day + temperature + (1 \| ID) | **0.146** | 0.048 | 0.285 |
| **Pair housing** |  |  |  |
| **Model** | **R** | **$CI_{low}$** | **$CI_{up}$** |
| Movement ~ time + (1 \| ID) | **0.041** | 0.015 | 0.067 |
| Shelter ~ time + (1\|ID) | **0.188** | 0.068 | 0.322 |
| Front ~ temperature + sex + time + (1 \| ID) | **0.119** | 0.049 | 0.182 |
| Heatmat ~ day + time + eggs presnet + temperature + (1 \| ID) | **0.289** | 0.112 | 0.503 |

## References

Alligood, C., & Leighty, K. (2015). Putting the “E” in SPIDER: Evolving trends in the evaluation of environmental enrichment efficacy in zoological settings. Animal Behavior and Cognition, 2(3), 200–217. 10.12966/abc.08.01.2015

Bashaw, M. J., Gibson, M. D., Schowe, D. M., & Kucher, A. S. (2016). Does enrichment improve reptile welfare? Leopard geckos (*Eublepharis macularius*) respond to five types of environmental enrichment. Applied Animal Behaviour Science, 184, 150–160. 10.1016/j.applanim.2016.08.003

Burghardt, G. M. (2013). Environmental enrichment and cognitive complexity in reptiles and amphibians: Concepts, review, and implications for captive populations. Applied Animal Behaviour Science, 147(3-4), 286–298. 10.1016/j.applanim.2013.04.013

Burghardt, G. M., Ward, B., & Rosscoe, R. (1996). Problem of reptile play: Environmental enrichment and play behavior in a captive Nile soft-shelled turtle, *Trionyx triunguis*. Zoo Biology, 15(3), 223–238. 10.1002/(SICI)1098-2361(1996)15:3<223::AID-ZOO3>3.0.CO;2-D

Burman, O. H. P., Collins, L. M., Hoehfurtner, T., Whitehead, M., & Wilkinson, A. (2016). Cold-blooded care: understanding reptile care and implications for their welfare. Testudo, 8(3), 83–86.

Bürkner, P.-C. (2017). brms: An R Package for Bayesian Multilevel Models Using Stan. Journal of Statistical Software, 80(1), 1–28. 10.18637/jss.v080.i01

Bürkner, P.-C. (2018). Advanced Bayesian multilevel modeling with the R Package brms. The R Journal, 10(1), 395–411. 10.48550/arXiv.1705.11123

Bürkner, P.-C. (2021). Bayesian Item Response Modeling in R with brms and Stan. Journal of Statistical Software, 100(5), 1–54. 10.48550/arXiv.1905.09501

Cabanac, M., & Bernieri, C. (2000). Behavioral rise in body temperature and tachycardia by handling of a turtle (*Clemmys insculpta*). Behavioural Processes, 49(2), 61–68. 10.1016/S0376-6357(00)00067-X

Cabanac, M., & Gosselin, F. (1993). Emotional fever in the lizard *Callopistes maculatus* (Teiidae). Animal Behaviour-London-Bailliere Tindall, 46, 200–200.

Damas-Moreira, I., Oliveira, D., Santos, J. L., Riley, J. L., Harris, D. J., & Whiting, M. J. (2018). Learning from others: an invasive lizard uses social information from both conspecifics and heterospecifics. Biology Letters, 14(10), 20180532. 10.1098/rsbl.2018.0532

Divers, S. J., & Mader, D. R. (Eds.). (2005). Reptile medicine and surgery-e-book. Elsevier Health Sciences.

Doody, J. S. (2023). Social Behaviour as a Challenge for Welfare. In Health and welfare of captive reptiles (pp. 189–209). Cham: Springer International Publishing.

Doody, J. S., Burghardt, G. M., & Dinets, V. (2013). Breaking the social–non-social dichotomy: a role for reptiles in vertebrate social behavior research? Ethology, 119(2), 95–103. 10.1111/eth.12047

Doody, J. S., Dinets, V., & Burghardt, G. M. (2021). The secret social lives of reptiles. JHU Press.

Duncan, I. J. (1998). Behavior and behavioral needs. Poultry Science, 77(12), 1766–1772. 10.1093/ps/77.12.1766

Estevez, I., & Christman, M. C. (2006). Analysis of the movement and use of space of animals in confinement: The effect of sampling effort. Applied Animal Behaviour Science, 97(2-4), 221–240. 10.1016/j.applanim.2005.01.013

Font, E., Burghardt, G. M., & Leal, M. (2023). Brains, Behaviour, and Cognition: Multiple Misconceptions. In Health and welfare of captive reptiles (pp. 211–238). Cham: Springer International Publishing.

Gainsbury, A. M. (2020). Influence of size, sex, and reproductive status on the thermal biology of endemic Florida scrub lizards. Ecology and Evolution, 10(23), 13080–13086. 10.1002/ece3.6897

Gillingham, J. C., & Clark, D. L. (2023). Normal Behaviour. In Health and welfare of captive reptiles (pp. 143–188). Cham: Springer International Publishing.

Grossmann, W. (2006). Der Tokeh, Gekko gecko. Münster: Natur und Tier Verlag.

Halliwell, B., Uller, T., Holland, B. R., & While, G. M. (2017). Live bearing promotes the evolution of sociality in reptiles. Nature Communications, 8(1), 2030. 10.1038/s41467-017-02220-w

Hurst, J. L., Barnard, C. J., Nevison, C. M., & West, C. D. (1997). Housing and welfare in laboratory rats: welfare implications of isolation and social contact among caged males. Animal Welfare, 6(4), 329–347. 10.1017/S0962728600020042

Hurst, J. L., Barnard, C. J., Nevison, C. M., & West, C. D. (1998). Housing and welfare in laboratory rats: the welfare implications of social isolation and social contact among females. Animal Welfare, 7(2), 121–136. 10.1017/S0962728600020455

Januszczak, I. S., Bryant, Z., Tapley, B., Gill, I., Harding, L., & Michaels, C. J. (2016). Is behavioural enrichment always a success? Comparing food presentation strategies in an insectivorous lizard (*Plica plica*). Applied Animal Behaviour Science, 183, 95–103. 10.1016/j.applanim.2016.07.009

Juri, G. L., Chiaraviglio, M., & Cardozo, G. (2018). Do female reproductive stage and phenotype influence thermal requirements in an oviparous lizard? Journal of Thermal Biology, 71, 202–208. 10.1016/j.jtherbio.2017.11.013

Langkilde, T., & Shine, R. (2006). How much stress do researchers inflict on their study animals? A case study using a scincid lizard, *Eulamprus heatwolei*. Journal of Experimental Biology, 209, 1035–1043. 10.1242/jeb.02112

Lenth R (2023). emmeans: Estimated Marginal Means, aka Least-Squares Means_. R package version 1.8.6–090001, <https://github.com/rvlenth/emmeans>.

Lillywhite, H. B. (2023). Physiology and functional anatomy. In Health and welfare of captive reptiles (pp. 7–44). Cham: Springer International Publishing.

Loew, E. R. (1994). A third, ultraviolet-sensitive, visual pigment in the Tokay gecko (*Gekko gecko*). Vision Research, 34(11), 1427–1431. 10.1016/0042-6989(94)90143-0

Meehan, C. L., Garner, J. P., & Mench, J. A. (2003). Isosexual pair housing improves the welfare of young Amazon parrots. Applied Animal Behaviour Science, 81(1), 73–88. 10.1016/S0168-1591(02)00238-1

Morgan, K. N., & Tromborg, C. T. (2007). Sources of stress in captivity. Applied Animal Behaviour Science, 102(3-4), 262–302. 10.1016/j.applanim.2006.05.032

Moszuti, S. A., Wilkinson, A., & Burman, O. H. (2017). Response to novelty as an indicator of reptile welfare. Applied Animal Behaviour Science, 193, 98–103. 10.1016/j.applanim.2017.03.018

Noble, D. W., Byrne, R. W., & Whiting, M. J. (2014). Age-dependent social learning in a lizard. Biology Letters, 10(7), 20140430. 10.1098/rsbl.2014.0430

Oka, T. (2018). Stress-induced hyperthermia and hypothermia. Handbook of clinical neurology, 157, 599–621. 10.1016/B978-0-444-64074-1.00035-5

Oufiero, C. E., & Angilletta Jr, M. J. (2010). Energetics of lizard embryos at fluctuating temperatures. Physiological and Biochemical Zoology, 83(5), 869–876.

PFMA, 2016. Pet Population Report. London, UK. 10.1086/656217

PFMA, 2022. Annual report (online) https://ukpetfood-reports.co.uk/annual-report-2022/

Preest, M. R., & Cree, A. (2008). Corticosterone treatment has subtle effects on thermoregulatory behavior and raises metabolic rate in the New Zealand common gecko, Hoplodactylus maculatus. Physiological and Biochemical Zoology, 81(5), 641–650. 10.1086/590371

R Core Team (2022). R: A language and environment for statistical computing. R Foundation for Statistical Computing, Vienna, Austria. URL https://www.R-project.org/. Accessed March 2023

Rey, S., Huntingford, F. A., Boltana, S., Vargas, R., Knowles, T. G., & Mackenzie, S. (2015). Fish can show emotional fever: stress-induced hyperthermia in zebrafish. Proceedings of the Royal Society B: Biological Sciences, 282(1819), 20152266. 10.1098/rspb.2015.2266

Robinson, J. E., St. John, F. A., Griffiths, R. A., & Roberts, D. L. (2015). Captive reptile mortality rates in the home and implications for the wildlife trade. PloS one, 10(11), e0141460. 10.1371/journal.pone.0157519

Rodríguez-Díaz, T., & Brana, F. (2011). Shift in thermal preferences of female oviparous common lizards during egg retention: insights into the evolution of reptilian viviparity. Evolutionary Biology, 38, 352–359. 10.1007/s11692-011-9122-y

Rodríguez-Díaz, T., González, F., Ji, X., & Braña, F. (2010). Effects of incubation temperature on hatchling phenotypes in an oviparous lizard with prolonged egg retention: are the two main hypotheses on the evolution of viviparity compatible? Zoology, 113(1), 33–38. 10.1016/j.zool.2009.05.001

Rosier, R. L., & Langkilde, T. (2011). Does environmental enrichment really matter? A case study using the eastern fence lizard, *Sceloporus undulatus*. Applied Animal Behaviour Science, 131(1-2), 71–76. 10.1016/j.applanim.2011.01.008

Stoffel, M. A., Nakagawa, S., & Schielzeth, H. (2017). rptR: repeatability estimation and variance decomposition by generalized linear mixed-effects models. Methods in Ecology and Evolution, 8, 1639–1644. 10.1111/2041-210X.12797

Sherwin, C. M., Christiansen, S. B., Duncan, I. J. H., Erhard, H. W., Lay, D. C., Mench, J. A., O’Connor, C. E., & Petherick, C. J. (2003). Guidelines for the ethical use of animals in applied animal behaviour research. Applied Animal Behaviour Science, 81, 291–305.

Shepherdson, D. (1998). Tracing the path of environmental enrichment in zoos. In Second nature: Environmental enrichment for captive animals (pp 1–12). Washington D.C: Smithsonian Institution.

Szabo, B., & Ringler, E. (2023). Fear of the new? Geckos hesitate to attack novel prey, feed near objects and enter a novel space. Animal Cognition, 26(2), 537–549.

Telemeco, R. S., Radder, R. S., Baird, T. A., & Shine, R. (2010). Thermal effects on reptile reproduction: adaptation and phenotypic plasticity in a montane lizard. Biological Journal of the Linnean Society, 100(3), 642–655. 10.1111/j.1095-8312.2010.01439.x

Tetzlaff, S. J., Sperry, J. H., & DeGregorio, B. A. (2022). You can go your own way: No evidence for social behavior based on kinship or familiarity in captive juvenile box turtles. Applied Animal Behaviour Science, 248, 105586. 10.1016/j.applanim.2022.105586

Troyer, K. (1987). Small differences in daytime body temperature affect digestion of natural food in a herbivorous lizard (*Iguana iguana*). Comparative Biochemistry and Physiology A, 87(3), 623–626.

Uller, T., While, G. M., Cadby, C. D., Harts, A., O’Connor, K., Pen, I., & Wapstra, E. (2011). Altitudinal divergence in maternal thermoregulatory behaviour may be driven by differences in selection on offspring survival in a viviparous lizard. Evolution, 65(8), 2313–2324. 10.1111/j.1558-5646.2011.01303.x

Visser, E. K., Ellis, A. D., & Van Reenen, C. G. (2008). The effect of two different housing conditions on the welfare of young horses stabled for the first time. Applied Animal Behaviour Science, 114(3-4), 521–533. 10.1016/j.applanim.2008.03.003

Vitt, L., Caldwell, J., Sartorius, S., Cooper, W., Baird, T., Baird, T., & Pérez-Mellado, V. (2005). Pushing the edge: extended activity as an alternative to risky body temperatures in a herbivorous teiid lizard (*Cnemidophorus murinus*: Squamata). Functional Ecology, 19, 152–158. 10.1111/J.0269-8463.2005.00947.X.

Wilkinson, A., Kuenstner, K., Mueller, J., & Huber, L. (2010). Social learning in a non-social reptile (*Geochelone carbonaria*). Biology Letters, 6(5), 614–616. 10.1098/rsbl.2010.0092

Williams, I., Hoppitt, W., & Grant, R. (2017). The effect of auditory enrichment, rearing method and social environment on the behavior of zoo-housed psittacines (Aves: Psittaciformes); implications for welfare. Applied Animal Behaviour Science, 186, 85–92. 10.1016/j.applanim.2016.10.013

Woolrich-Piña, G. A., Smith, G. R., Lemos-Espinal, J. A., & Ramírez-Silva, J. P. (2015). Do gravid female Anolis nebulosus thermoregulate differently than males and non-gravid females? Journal of Thermal Biology, 52, 84–89. 10.1016/j.jtherbio.2015.06.006

Yan, X. F., Tang, X. L., Yue, F., Zhang, D. J., Xin, Y., Wang, C., & Chen, Q. (2011). Influence of ambient temperature on maternal thermoregulation and neonate phenotypes in a viviparous lizard, *Eremias multiocellata*, during the gestation period. Journal of Thermal Biology, 36(3), 187–192. 10.1016/j.jtherbio.2011.02.005

